# Emergence of Anticipatory Beta Activity to Facilitate Behavioral Stability Following Environmental Changes

**DOI:** 10.1101/2024.12.23.630078

**Authors:** Martina Bracco, Varsha Vasudevan, Vridhi Rohira, Quentin Welniarz, Mihoby Razafinimanana, Alienor Richard, Christophe Gitton, Sabine Meunier, Antoni Valero-Cabré, Denis Schwartz, Traian Popa, Cécile Gallea

## Abstract

Adaptive behavior enables flexible responses to environmental changes. This process is particularly crucial when transitioning between environments with different features, relying on the progressive formation of expectations based on prior experience. In humans, beta oscillations are central to adaptive behavior. Yet, the brain mechanisms underlying the detection of environmental changes, and the iterative update needed to progressively improve behavioral performance remain elusive. Here, we reveal that beta activity emerges in a cerebello-cortical network two seconds before action initiation, as the features of a new environment become known and behavioral outcomes become more predictable. Within this period, the cerebellum and parietal cortex drive prefrontal activity to form expectations. Using a single-trial approach, we establish that beta bursts before action initiation predict performance in the upcoming trial based on previous outcomes. These findings uncover a novel anticipatory mechanism that reflects predictive processes critical for stabilizing performance and adapting to environmental changes.

## Main

Successful adaptation to different environments depends on detecting their features and rapidly adjusting behavior to maintain optimal performance. This is the case of a tennis player recalibrating their forehand to counteract a headwind. This dynamic process relies on predictive mechanisms, where *outcome expectations*, shaped by prior experience, are continuously compared to following outcomes. Any mismatch generates *prediction errors*, which arise when action feedback becomes available and contradicts these expectations. Prediction errors are largest as the brain confronts unfamiliar environmental features, but they diminish as learning progresses and a more accurate representation of the environment emerges. Trial-by-trial, the brain integrates past errors to update its expectations, progressively forming a more predictable representation of the environment (*1*). Therefore, the brain and behavior continuously oscillate between *stable* states, characterized by small behavioral fluctuations, and *transition* states, marked by large fluctuations in response to environmental changes that progressively recover to establish a new stable state. In this framework, brain and behavior states *emerge* from the interaction between the body’s biomechanics, environmental constraints, and task goal (*2*). But how does the brain process state-dependent information to transition from an unknown to a stable and predictable environment, hence enabling the prediction of future outcomes?

Beta oscillations (13-35Hz) play a crucial role in adaptative processes (*3–9*). In sensorimotor adaptation, pre-movement beta synchronization is modulated by prior errors (*10*– *12*). Up to two seconds before movement onset, beta synchronization decreases following erroneous movements caused by unexpected perturbations, and increases after successful movements (*10*, *11*). These findings suggest that pre-movement beta oscillations encode expectations based on past performance (*13*, *14*). Yet, in these studies, the perturbations were transient, so that past trials cannot be used to update expectations and improve future outcomes adequately. Moreover, the spatial resolution EEG employed in previous studies constrains our understanding of the specific neural circuits and regions involved in generating beta oscillations, leaving open questions about their network-level integration. Indeed, beta oscillations are ubiquitous in the brain and have been identified in several areas, including the motor, parietal, and frontal cortices, as well as the cerebellum and the basal ganglia (*15–25*). Such a widespread distribution may imply that beta oscillations support the flow of expectations across brain areas (*26*), participating in progressive motor adjustments in new environments with different features.

The cerebellum and the parietal cortex have been identified as critical in predictive processes during motor adaptation (*27*). The cerebellum acts as an online ‘comparator’ between the motor plan and the actual sensory feedback, to adjust movements during the execution phase (*28–30*). Furthermore, recent work has shown that cerebellar activity increases well before movement execution (*31–33*), suggesting an involvement in anticipatory processes, such as motor preparation and planning (*34*, *35*). Like the cerebellum, the parietal cortex is active at the early stages of motor planning, providing information to compensate for motor and target errors (*19*, *36*). The parietal cortex acts as an integrator of somatosensory information and maintains predictions of the sensory consequences of movements (*37*). Additionally, it supplies the prefrontal cortex with essential contextual information for goal-generation (*38*). This suggests that the parietal cortex, and possibly the cerebellum, may communicate expectations to frontal and prefrontal areas through anticipatory signals used to update motor plans before movement initiation (*26*, *39*, *40*). Such signals may allow the brain to prepare for action based on experience, as when an expert tennis player anticipates the outcome of a serve based on subtle environmental cues, often before the ball has been hit.

Here, we aimed to characterize the brain network dynamics that mediate adaptive behavior through pre-movement beta oscillations. Specifically, we sought to determine how outcome expectations and the emergence of specific brain activity patterns occur over time to facilitate motor performance. To this aim, we recorded magnetoencephalography (MEG) from healthy human participants as they performed blocks of 50 consecutive trials of a reaching task under different environmental characteristics; i.e., either normal (no rotation) or rotated visual feedback (rotation). Using a joystick, participants controlled the position of a cursor on a screen. In ‘normal’ visual feedback blocks, the cursor’s trajectory matched the joystick position, resulting in minimal trajectory errors and stable movement outcomes. In contrast, rotated visual feedback introduced a perceptual bias, leading to significant trajectory errors during early trials. These errors decreased and stabilized over the 50 trials as participants adapted to the perturbation (*41*, *42*).

We employed two complementary approaches to answer our questions. Using a conventional averaging approach, we first identified the effect of different states (transition *vs.* stable) and environmental perturbations (no rotation *vs.* rotation) on the spatiotemporal features of pre-movement beta oscillations to identify a network of brain areas (cluster analysis) and on the information flow within this network (effective connectivity analysis) in a time window preceding the target appearance, i.e. before a precise movement plan had occurred. While these conventional analyses provide insight into the functional role that a specific network may play, they do not reveal how specific processes unfold through time, nor do they clarify how the coordinated interactions among a network of brain regions translate into behavior (*43*). To account for this important point, we then employed a single-trial approach to determine the trial-by-trial relationship between pre-movement beta activity and behavior. This is particularly important for adaptative behavior, where trial-by-trial variations of beta activity could reveal important dynamics as individual trials carry critical information for progressively improving behavioral performance. Furthermore, single-trial analyses have recently shown beta oscillations within cortical and subcortical brain areas emerge as brief events or bursts during the pre-movement period (*22*, *44–48*). For example, in single trials, beta bursts before the onset of a simple movement correlate with the kinematics during its execution (*44*) and predict movement initiation timing in decision tasks guided by visual cues (*45*, *48*). On such a basis, we used a predictive model that simultaneously captured the influence of previous errors on current motor performance and the effect of beta bursts on subsequent motor performance, providing new insights into the neural mechanisms of adaptive behavior.

## Results

### Anticipatory beta activity supports the update and maintenance of outcome expectations

#### Behavior

Participants performed blocks of 50 consecutive trials of a reaching task with different environmental characteristics, *i.e.*, either with normal (0°; no rotation) or rotated visual feedback (25°, −30°; rotation; Figure 1). These blocks were randomly interleaved across participants, inducing a transition from one environment to another every 50 trials. This transition ensured that while participants’ performance stabilized within a block, the alternation between different environments hindered their ability to consolidate adaptive performance across blocks (*43*). This distinction is crucial for dissociating the effect of transition between environments from simple motor adjustments over trials.

**Fig. 1.**
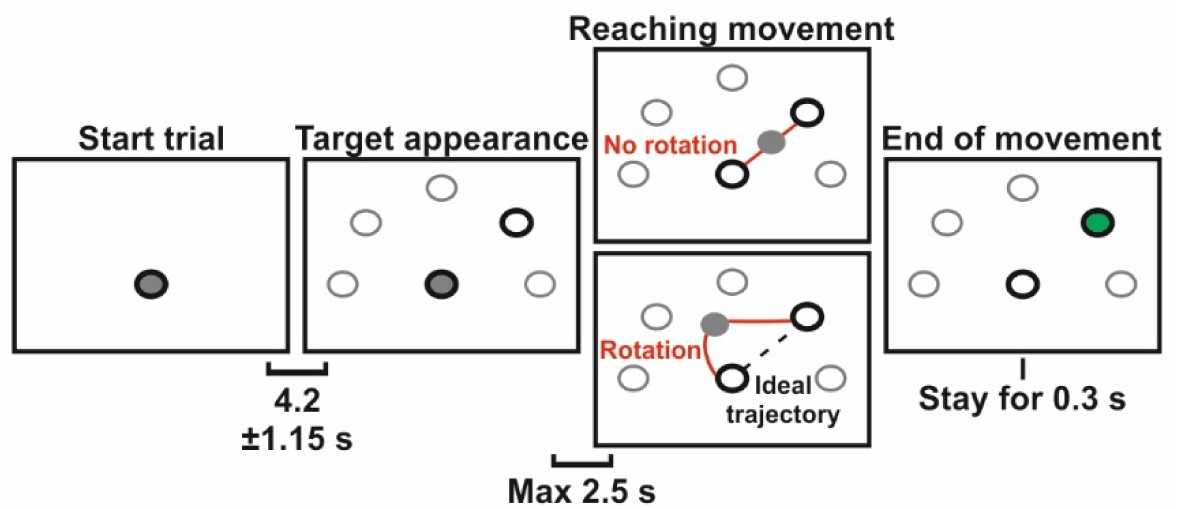
Reaching task description. Each trial began with the presentation of a solid gray cursor within a fixation circle at the bottom center of the screen. After 4.2 s (jittered by ±1.15 s), a circular black target appeared at one of five possible locations. We instructed participants to swiftly move the cursor toward the visual target, aiming to follow the straightest path (ideal trajectory). Each reaching movement had a maximum duration of 2.5 s; if this limit was exceeded, the target disappeared, and participants were prompted to return the joystick to the central position. A trial ended when the cursor remained within the target for 0.3 s, causing the target to turn green and disappear. Participants then returned the cursor to the central fixation circle. Participants performed blocks of consecutive trials of the same perturbation. In the “No rotation” blocks, the cursor’s trajectory aligned with the joystick movement and expected visual feedback. In the “Rotation” blocks, the visual feedback of the trajectory was rotated either 25° clockwise (25° rotation) or 30° counterclockwise (−30° rotation) relative to the real trajectory. The red line indicates an example of trajectory during the transition state of both “No rotation” and “Rotation” blocks, while the black dashed line represents the ideal trajectory.

Participants controlled a cursor using a joystick with the right dominant hand. The goal of the task was to reach a peripheral target, as fast as possible, starting from a central position. Participants’ motor performance was estimated as trajectory errors on each trial. This was quantified as the area under the curve (AUC) representing the cursor’s trajectory deviating from an ideal straight line between the starting and the final target position (higher AUC values indicate greater errors). To characterize changes in participants’ performance, we divided each block with the same environmental characteristics (*Perturbation*) into two *States*: a transition state (first 16 trials), in which participants showed an overall 80% performance improvement, and a stable state (last 16 trials) in which performance stabilized (figure 2A). This division was based on the observed population’s errors within blocks with rotated feedback (see Methods for details). As expected, errors decreased significantly from the transition to the stable state (2×2 repeated measures ANOVA; Main Effect of *State* (transition *vs.* stable): F(1,16) = 560.27, p < 0.001; η_p_^2^ = 0.97). However, participants consistently made more errors in blocks with rotation compared to those with no rotation (2×2 repeated measures ANOVA; Main Effect of *Perturbation* (rotation *vs.* no rotation): F(1,16) = 155.56, p < 0.001; η_p_^2^ = 0.67). This difference persisted across all states, with substantially more errors in stable ones orotation *vs.* rotation, transition state: t = −15.66, p < 0.001, Cohen’s d = −4.57; no rotation *vs.* rotation, stable state: t = −7.13, p < 0.001, Cohen’s d = −2.08). Notably, the reduction in errors was mainly observed in the environment when rotation was applied (2×2 repeated measures ANOVA; *State* x *Perturbation* Interaction: F(1,16) = 110.536, p < 0.001; η_p_^2^ = 0.09; transition *vs.* stable, no rotation: t = 2.28, p = 0.19, Cohen’s d = 0.31; transition *vs.* stable, rotation: t = 20.67, p < 0.001, Cohen’s d = 2.80; Figure 2B).

**Fig. 2.**
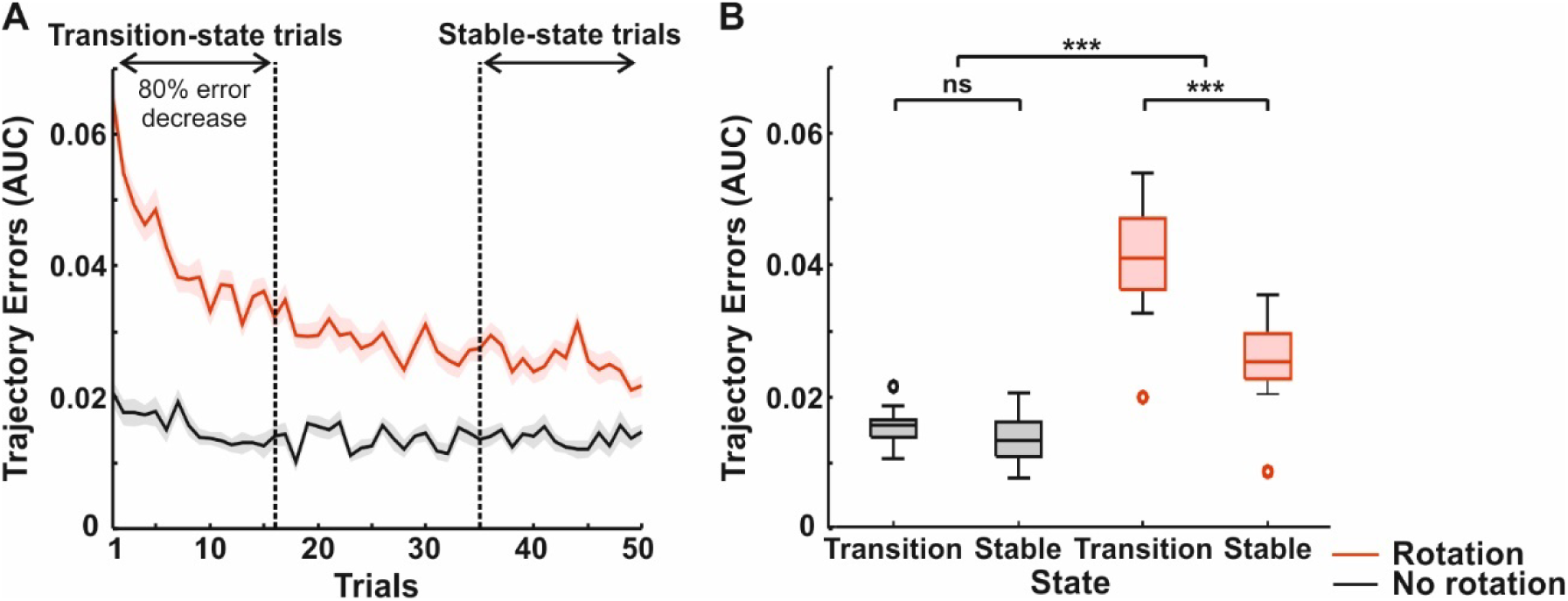
Behavioral performance results. (A) Evolution of trajectory error amplitude (AUC) along 50 consecutive trials of a block, averaged across participants relative to *Perturbation* (Rotation, red; No rotation black). Shading indicates SEM. Vertical dashed lines divide the blocks into transition (first 16 trials) and stable state(last 16 trials). (B) Results of the 2×2 repeated measures ANOVA comparing the trajectory errors categorized by *State* (transition *vs.* stable) and *Perturbation* (rotation *vs.* no rotation). Box plots show the median (horizontal line), the lower and upper quartiles (box), the minimum and maximum values that are not outliers (whiskers), and the outliers, computed using the interquartile range (dots). *** indicates p < 0.001; ns indicates p > 0.05.

### Averaged pre-movement beta oscillatory activity (MEG): sensor and source space

We recorded participants’ brain activity during the reaching task to determine the spatiotemporal changes of pre-movement beta synchronization (14-26 Hz) across different states and environmental perturbations. To achieve this, for each participant, we transformed the MEG data into a time-frequency domain, averaging across trials to obtain separate event-related, average, power spectra corresponding to different *States* (transition and stable) and *Perturbations* (rotation and no rotation; Figure 3), as we did for the behavioral analysis. Individual data were entered into a 2×2 non-parametric cluster-based approach and paired t-tests.

**Fig. 3.**
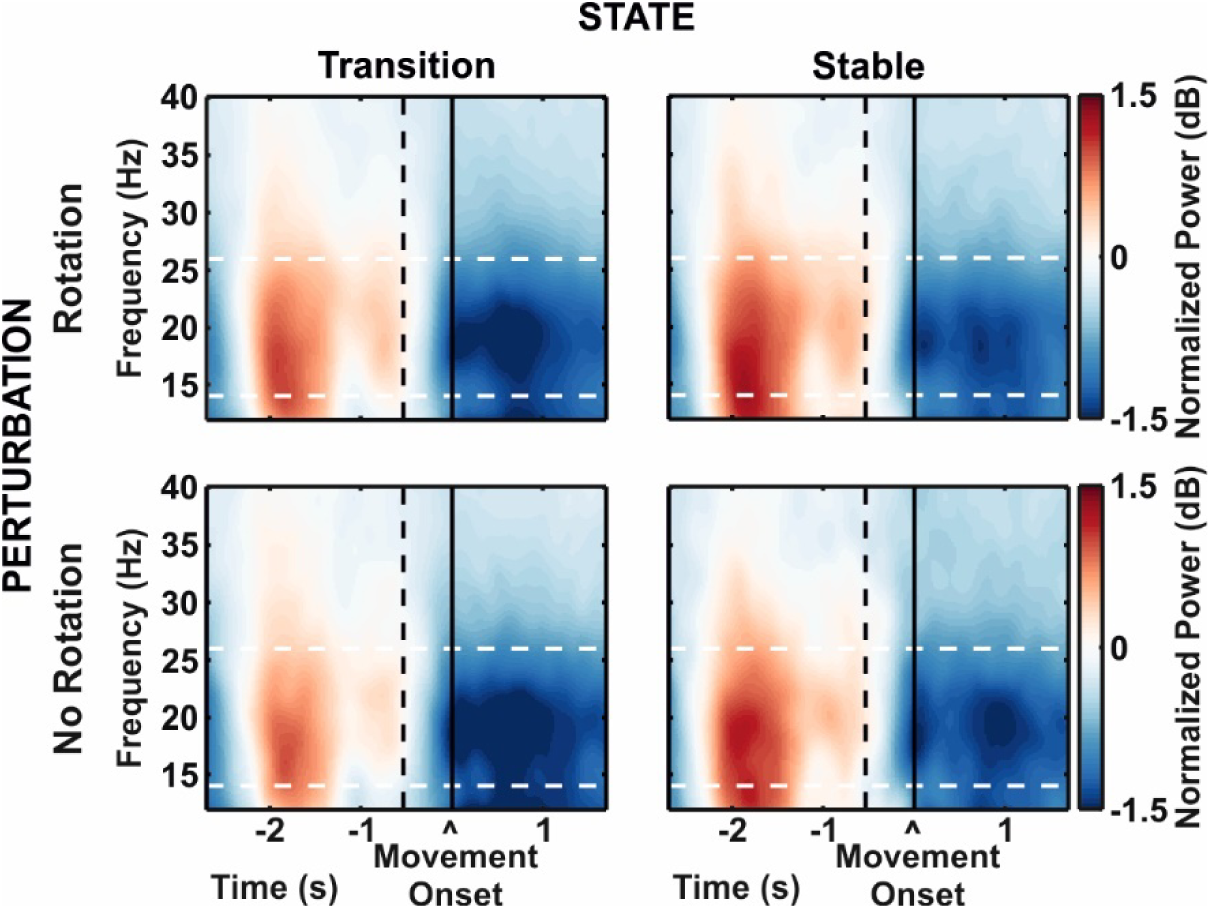
Time-frequency power spectra across different States and Perturbations. The time-frequency plots illustrate the power spectrum for frequencies ranging from 12 to 40 Hz, spanning −2.5 to 0 seconds relative to movement onset. Power was normalized by dividing each time-frequency-sensor value by the average power across all trials, independent of behavioral conditions, which was then log-transformed. For visualization, data was averaged across all 204 horizontal gradiometers and all participants. The black vertical dashed line indicates the median target appearance (i.e., −0.5 s); the black vertical solid line indicates the movement onset (*i.e.*, 0 s); the white horizontal dashed lines highlight the beta frequency range analyzed (14-26 Hz).

We found a significant overall increase in pre-movement beta synchronization from transition to stable states (2.5-1.35 s before movement onset; cluster analysis; Main Effect of *State* (transition *vs.* stable): p < 0.001) in both perturbation conditions (rotation (transition *vs*. stable): p = 0.009; no rotation (transition *vs.* stable): p = 0.003; Figure 4A). Source localization revealed that this change primarily originated from the cerebellum and temporal cortices bilaterally (Figure 4A, Table S1). Interestingly, when passing from a transition to a stable state, pre-movement beta synchronization showed a smaller increase during rotation than in no rotation blocks (2.35-2 s before movement onset; cluster analysis; *State* x *Perturbation* Interaction: p = 0.014; Figure 4B). This interaction between states and perturbations was not attributable to a difference in beta power during the transition state (cluster analysis; transition (rotation *vs*. no rotation); all p values > 0.229) or to an overall difference across perturbations (rotation *vs.* no rotation; all p values > 0.073). The source estimates revealed that this interaction originated from a network of regions including the left parietal cortex, the right frontal and prefrontal cortices, and the right cerebellum (Figure 4B, Table S1).

**Fig. 4.**
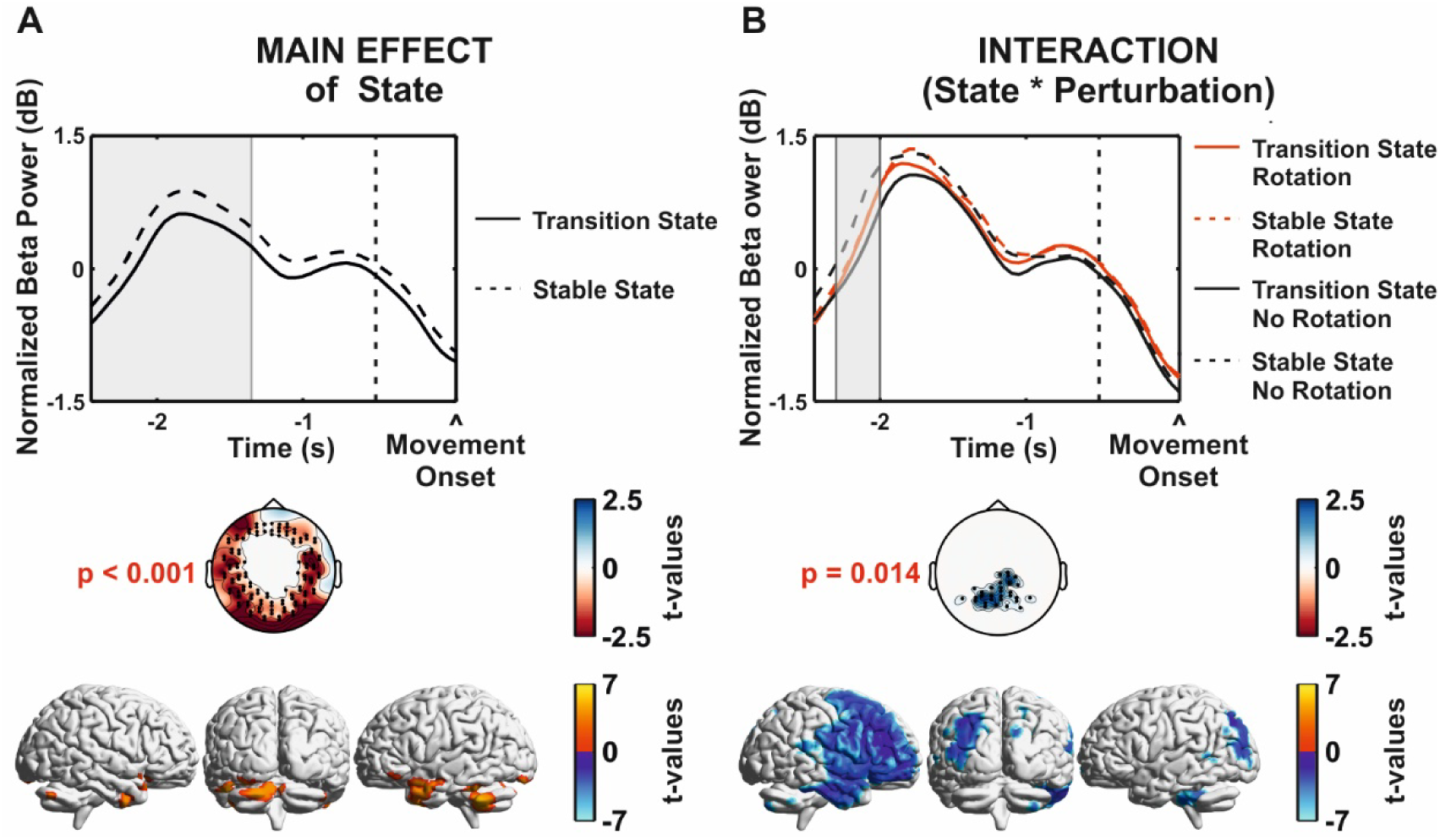
Averaged pre-movement beta oscillatory activity results. (**A**) Main effect of *State*, obtained by combining the data from both *Perturbations* (rotation and no rotation). (**B**) Interaction effect between *State* and *Perturbation*, obtained by comparing the difference in beta activity between transition and stable states within each perturbation ( (table, transition)). For each effect, the results of the cluster-based analysis are represented wit^Δ^h power spectra plots (top row), scalp topographies (central row), and source space representations (bottom row). For the power spectra plots, the two-dimensional clusters (time and sensors) are collapsed across the significant sensors (shown within the topography plots) to display the temporal evolution of the analyzed beta power (14-26 Hz). The vertical dashed line indicates the median target appearance (at −0.5 s), and shaded areas mark time windows significant in the cluster-based analysis (Main effect of *State*: −2.5 to −1.35 seconds; Interaction of *State* × *Perturbation*: −2.35 to −2 seconds). For scalp topographies, data are collapsed across significant time points to highlight spatial distributions. Only significant t-values are shown, and black dots indicate significant sensors (threshold: p < 0.05). The source space results (threshold: p < 0.001) are overlaid on a 3D brain surface template (bottom panel).

### Dynamics of pre-movement beta connectivity (MEG)

We performed an effective dynamic connectivity analysis in the beta range (14-26 Hz) during the 2.5 s preceding movement initiation. We computed connectivity using nonparametric conditional Granger causality during the transition and stable states, across both perturbations (rotation, no rotation). We specifically focused on those brain areas highlighted in our previous source space analysis and known to be relevant for sensorimotor adaptation and predictive processes: the posterior right cerebellum (rCB), the left parietal cortex (lPC), and the right prefrontal cortex (rPFC; (*35*)). To assess the temporal dynamics of such effective connectivity patterns, we examined five overlapping 0.45 s time windows before target appearance (from −2.5 to −2.05 s; from −2.15 to −1.7 s; from −1.8 to −1.35 s; from −1.45 to −0.9 s; from −1.0 to −0.65 s), capturing at least six cycles of beta oscillation at its lowest frequency (14 Hz).

Our results showed that, like beta power, connectivity increased from transition to stable states within the same block. This increase was reciprocal among the right cerebellum, the left parietal cortex, and the right prefrontal cortex (5×2×2 ANOVA (*Time Window x State x Perturbation*), Main Effect of *State*: Cerebellum to Prefrontal, F(1,16) = 6.57, p = 0.02, 0.29; Cerebellum to Parietal, F(1,16) = 3.88, p = 0.06, η_p_^2^ = 0.19; Parietal to Cerebellum, F(1,16) = 5.53, p = 0.03, η_p_^2^ = 0.26; Parietal to Prefrontal, F(1,16) = 11.97, p = 0.003, η_p_^2^ = 0.43; Prefrontal to Cerebellum, F(1,16) = 6.61, p = 0.02, η_p_^2^ = 0.29; Prefrontal to Parietal, F(1,16) = 5.36, p = 0.03, η_p_^2^ = 0.25; Figure 5A). However, an interesting pattern emerged in the five time windows considered in the pre-movement period. As the time window approached the movement initiation, connectivity specifically increased from both the cerebellum (5×2×2 ANOVA, Main Effect of *Time Window*: F(2.28,36.54) = 4.32, p = 0.02, η_p_^2^ = 0.20, Huynh-Feldt sphericity correction; −2.5/-2.05 *vs.* −1.45/-0.9: t = −3.51, p = 0.008, Cohen’s d = −0.36; −2.5/-2.05 *vs.* −1.0/- 0.65: t = −3.03, p = 0.04, Cohen’s d = −0.31; Figure 5B) and the parietal cortex (5×2×2 ANOVA, Main Effect of *Time Window*: F(2.65,42.36) = 3.16, p = 0.04, η_p_^2^ = 0.16, Huynh-Feldt sphericity correction; −2.5/-2.05 *vs.* −1.0/-0.65: t = −3.07, p = 0.02, Cohen’s d = −0.31; Figure 5B) to the prefrontal cortex. Notably, this increase was not bidirectional (5×2×2 ANOVA, Main Effect of *Time Window*, Prefrontal to Cerebellum: F(2.58,41.35) = 1.15, p = 0.34, η_p_^2^ = 0.07; Prefrontal to Parietal: F(1.55,24.75) = 0.87, p = 0.40, η_p_^2^ = 0.05). There was no significant effect of *Perturbation* (5×2×2 ANOVA, Main Effect of *Perturbation* Cerebellum to Prefrontal: F(1,16) = 2.21, p = 0.16, η_p_^2^ = 0.12; Parietal to Prefrontal: F(1,16) = 0.86, p = 0.37, η_p_^2^ = 0.05), indicating that the increase in connectivity from cerebellar and parietal areas to the prefrontal cortex was independent of the error magnitude.

**Fig. 5.**
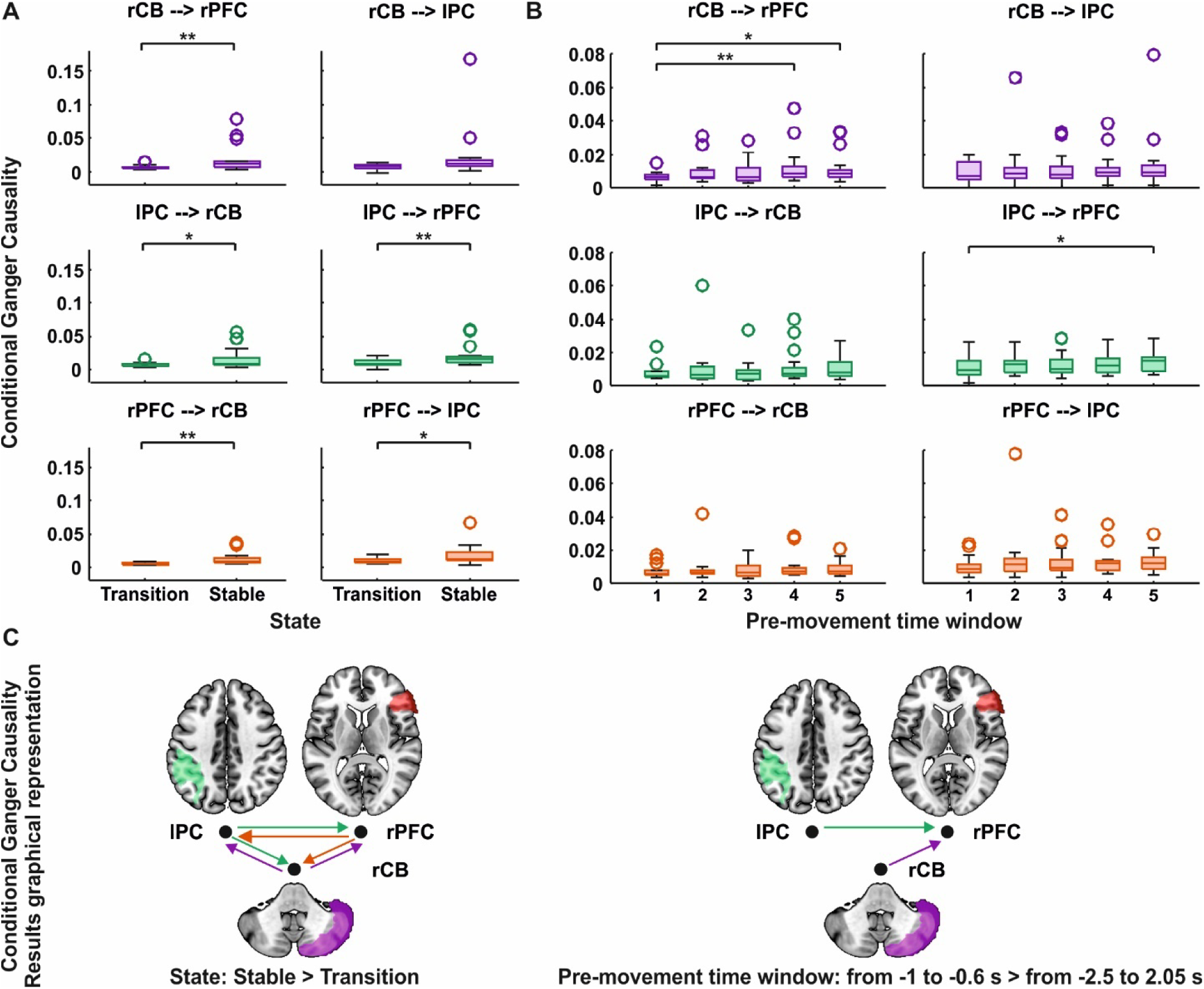
Conditional Granger causality results. (**A**) Main effect of *State* for the 5×2×2 repeated measures ANOVA (comparing transition *vs.* stable states independent of the other factors). (**B)** Main effect of *Time Window* from the same ANOVA, comparing the five time-windows (−2.5/-2.05; −2.15/-1.7; −1.8/-1.35; −1.4/-0.9; −1.0/- 0.65). (**C**) Graphical representation of the Conditional Granger Causality analysis showing the effect of *State* (left) and *Pre-movement window* (right). rCB: right cerebellum (Crus II; purple); rPFC: right prefrontal cortex (frontal inferior triangularis; organge); lPC: left parietal cortex (parietal inferior, green). Box plots show the median (horizontal line), the lower and upper quartiles (box), the minimum and maximum values that are not outliers (whiskers), and the outliers, computed using the interquartile range (dots). Dashed arrows reflect the connectivity direction. ** indicates p < 0.025; * indicates p < 0.05.

### Trial-by-trial beta burst dynamics encode expectations of future outcomes and support iterative motor improvement

#### Beta burst characterization

We sought to determine the trial-by-trial relationship between pre-movement beta bursts and trajectory errors in our task. To this aim, we isolated beta bursts in each trial of each block (Figure 6A). Beta bursts were defined as local maxima power exceeding a set power cut-off in the 2.5-second pre-movement epoch, at frequencies ranging from 14 to 26 Hz, and from sensors included in the significant clusters from the average beta power analysis (*State* x *Perturbation* Interaction; sensors shown in Figure 4B). The power cut-off was fixed to 3 times the median power, which provided the strongest correlation between the average mean power and the area covered by beta bursts (3X; all r values > 0.96; all p values < 0.001 Figure 6B), thus best representing the average beta power. This cut-off allowed us to detect ∼7 bursts per trial, each lasting less than 150 ms (Figure 6C).

**Fig. 6.**
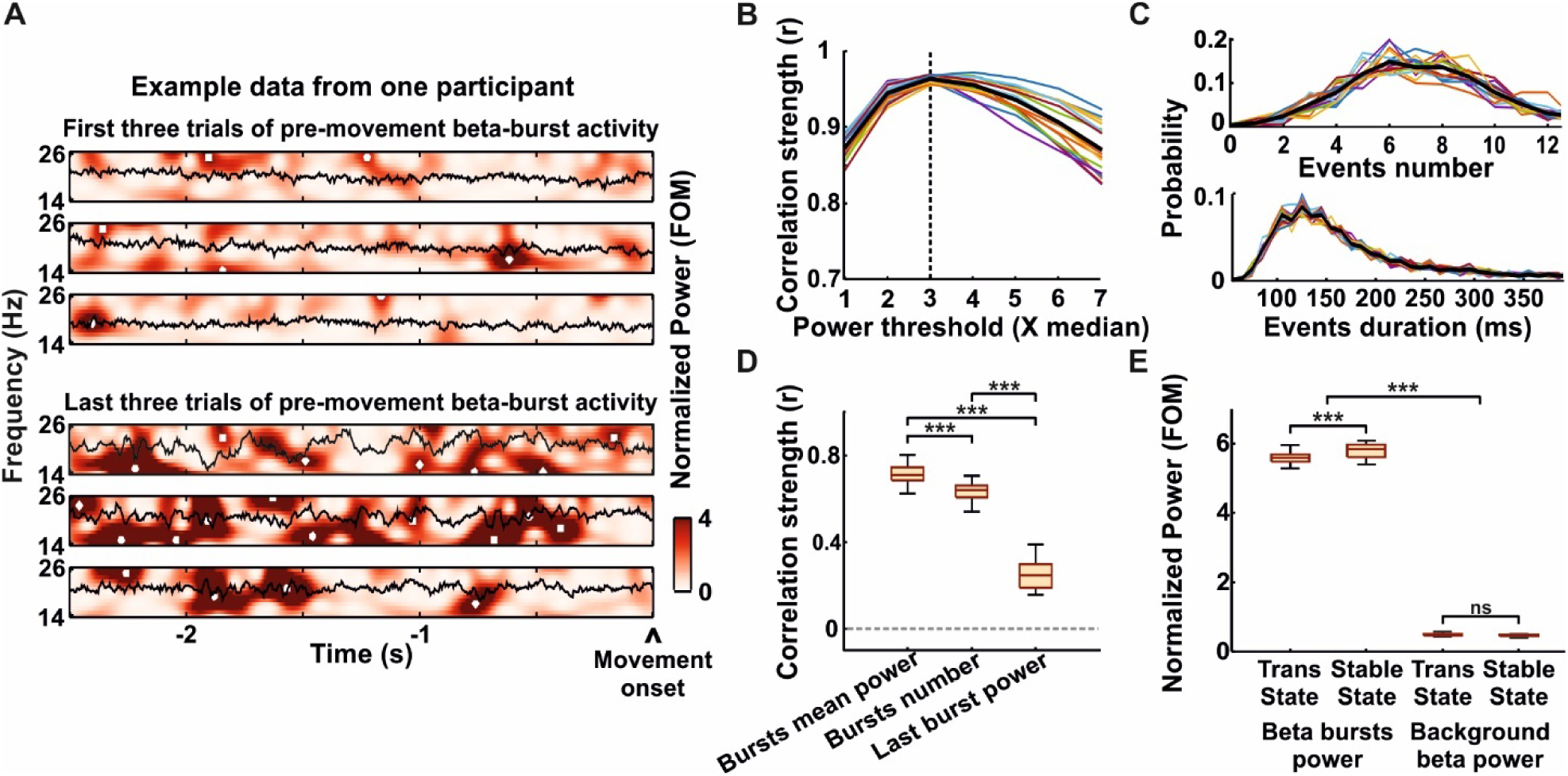
Beta bursts features. (**A**) Beta bursts in the 14–26 Hz range are shown from −2.5 seconds to movement onset for the first three and last three trials of an example block in a representative participant. Data were normalized as factors of the median (FOM) for each frequency, calculated separately for each participant and condition. (**B**) The threshold for data normalization was selected based on the highest correlation between the percentage of the spectrogram area exceeding thresholds ranging from 1 to 7 times the median power and the mean beta power for each trial. Colored lines represent the averages of trials and blocks for individual participants, while the black line shows the group average. The dashed vertical black line indicates the threshold with the highest correlation (3X median power). (**C**) Probability distribution of events number (top) and events duration (bottom), where an event is a beta burst exceeding the 3X median cut-off. Distributions are plotted as the average across trials and blocks for individual participants (colored lines), with the population average in black. (**D**) Correlations between selected beta burst features (mean beta burst power, number of bursts, and power of the last burst) and average beta power. (**E**) Control analysis results verifying the relevance of the selected beta feature (mean burst power) to error reduction. Box plots depict the results of the 2×2×2 repeated measures ANOVA, with *Feature* (beta burst power *vs.* background beta activity), *State* (transition *vs.* stable), and *Perturbation* (rotation *vs.* no rotation) as factors. Box plots show the median (horizontal line), the lower and upper quartiles (box), the minimum and maximum values that are not outliers (whiskers), and the outliers, computed using the interquartile range (dots). *** indicate p < 0.001.

Among potentially quantifiable beta features relevant to behavior (*i.e.*, bursts mean power, burst number, last burst power; (*49*)), the mean power of beta bursts showed the highest correlation with the average beta power (repeated measures ANOVA; Main effect of Beta *Feature* (beta bursts mean power *vs*. bursts number *vs*. last burst power): F(2,32) = 481.30, p < 0.001; p = 0.97; bursts mean power *vs*. bursts number: t = 5.04, p < 0.001, Cohen’s d = 1.54; bursts mean power *vs*. last bursts power: t = 29.03, p < 0.001, Cohen’s d = 8.84; bursts number *vs*. last bursts power: t = 23.99, p < 0.001, Cohen’s d = 7.31; Figure 6D). Additionally, the power of beta bursts contributed significantly to error reduction along the different states (transition *vs.* stable), compared to background beta activity (3×2×2 repeated measures ANOVA; *Feature* x *States* interaction: F(1,16) = 13.14, p = 0.002, η_p_^2^ = 0.45; Figure 6E). More specifically, the power of the beta bursts increased across states (beta bursts mean power; transition *vs*. stable state: t = −4.96, p < 0.001, Cohen’s d = −1.53), reflecting the average beta power trends, while the background activity remained stable (background beta activity; transition *vs*. stable state: t = 0.48, p = 1.00, Cohen’s d = 0.11). Altogether these results support the use of beta burst power as a potentially relevant variable to model the relationship between beta activity and motor adjustments over trials.

#### Trial-by-trial relationship between beta bursts and trajectory errors: predictive model

To investigate whether beta burst dynamics during the pre-movement period could predict trajectory errors across consecutive trials, we used a time-series modeling approach. Specifically, we employed a regression framework, the Auto Regressive Moving Average (regARMA), modeling the dynamics of trajectory errors over consecutive trials, and including the trial-wise beta burst power as an external predictor. The optimal model was selected based on the lowest Akaike Information Criteria (AIC) values, tested across all feasible model combinations on population-averaged data from the 17 participants, as detailed in the Methods section. The regARMA (1,1) model with a Gaussian distribution and no intercept was identified as the best-fitting model for all perturbations (Table S2).

We validated the selected regARMA (1,1) model using a leave-one-out prediction approach. For each participant, model parameters were estimated using the average data from the other 16 participants. These parameters were then used to predict the left-out participant’s error evolution across trials. Specifically, participant’s error in the first trial and the trial-by-trial beta burst evolution over 50 trials served to predict the error across the remaining 49 trials (Figures 7A; see Figures S1, S2, and S3 for individual simulations). This enabled the assessment of the model’s ability to generalize and accurately capture the relationship between beta burst evolution and motor performance across individual participants.

**Fig. 7.**
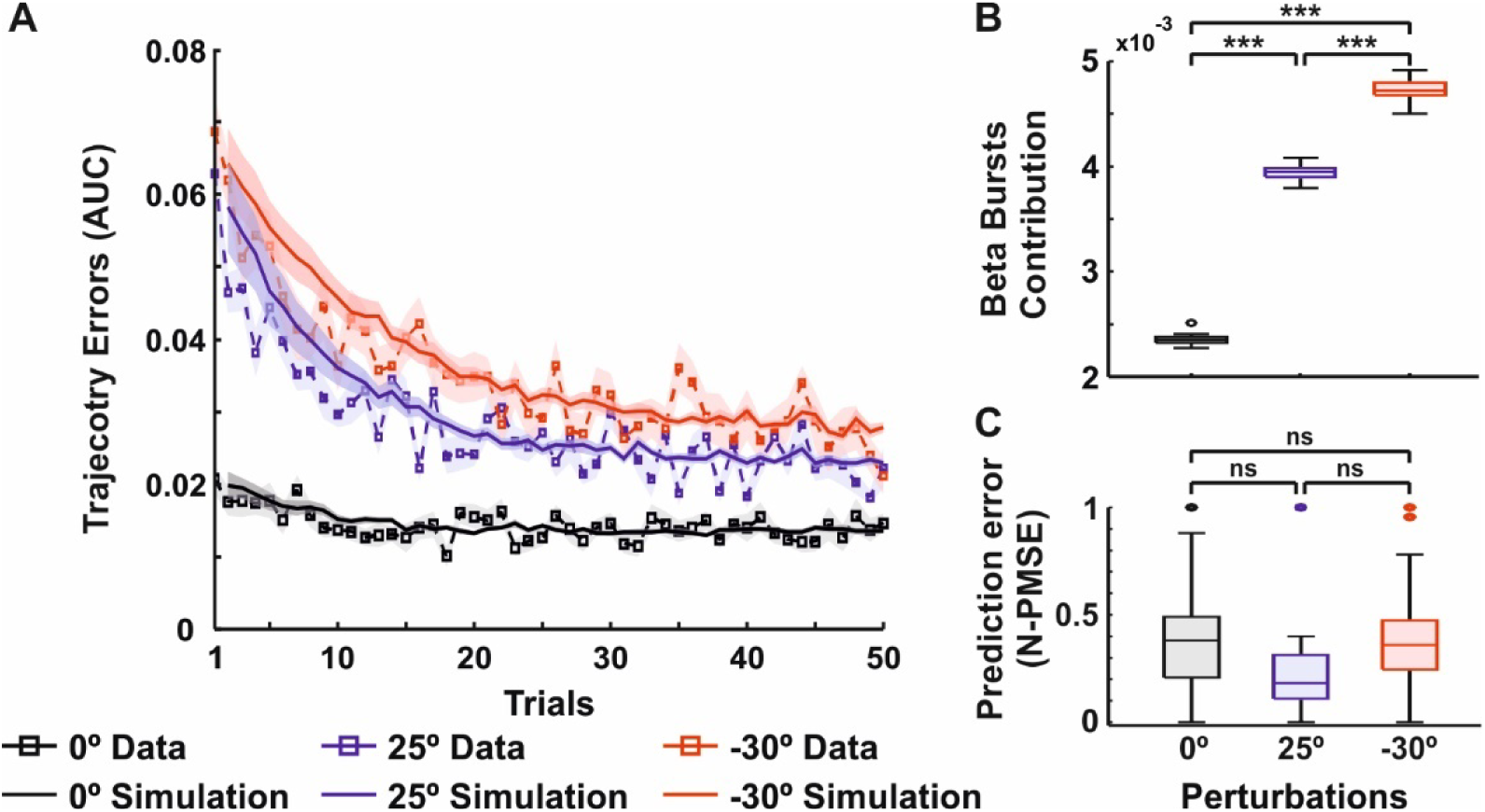
Predictive model results. (**A**) Real and predicted trajectory error amplitude along 50 consecutive trials of a block, averaged across participants for each perturbation (Rotation with −30° in red and 25° in blue; No rotation, black). The amplitude of real trajectory errors is quantified as the area under the curve (AUC), representing the deviation from the ideal straight-line trajectory between the start and target positions. Higher AUC values indicate greater trajectory error amplitude. Predicted errors are estimated by validating the chosen regARMA(1,1) model using a leave-one-out cross-validation approach. Shading indicates SEM. (**B**) Results of a repeated measures ANOVA comparing the coefficient of the regARMA model’s external predictor, representing the contribution of beta burst power to trajectory error evolution, across perturbations (0° *vs.* 25° *vs.* −30°). (**C**) Results of a repeated measures ANOVA comparing the normalized prediction mean squared error (N-PMSE) of the regARMA models across different perturbations (0° *vs.* 25° *vs.* −30°). Box plots show the median (horizontal line), the lower and upper quartiles (box), the minimum and maximum values that are not outliers (whiskers), and the outliers, computed using the interquartile range (dots). *** indicates p < 0.001; ns indicates p > 0.05.

We then used the coefficient of the regARMA model’s external predictor as a measure of beta burst power contribution to the motor adjustments over trials. Beta contribution defines the strength of the influence that pre-movement beta bursts have on motor performance, with higher values indicating a stronger role of beta oscillations in facilitating motor performance and reducing errors across trials. Our analysis revealed that the contribution of beta burst power varied across perturbations (repeated measures ANOVA, Main Effect of *Perturbation* (0° *vs.* 25° *vs.* − 30°): F(1.4,22.55) = 6244.68, p < 0.001, ^2^= 0.99, Huynh-Feldt sphericity correction; Figure 7B).

Specifically, the beta contribution incred with the increase of degrees of rotation (0° *vs.* 25°: t = −73.55, p < 0.001, Cohen’s d = −20.20; 0° *vs.* 30°: t = −109.64, p < 0.001, Cohen’s d = −30.10; 25° *vs.* 30°: t = −36.09, p < 0.001, Cohen’s d = −9.91; Figure 7B). Importantly, the model performed consistently across all perturbations, as indicated by non-significant differences in model prediction error (normalized mean squared error, N-PMSE; repeated measures ANOVA; Main Effect of *Perturbation* (0° *vs.* 25° *vs.* −30°): F (2,32) = 2.50, p = 0.10, η^2^= 0.14; Figure 7C). This suggests that the difference in beta contribution between different perturbations was not due to overall differences in the models’ generalizability.

#### Model specificity

We also evaluated whether the relationship between MEG activity and motor performance was specific to the beta frequency. To this aim, we tested the same regARMA model with an external predictor derived from an alternative frequency. We specifically selected alpha (8-12 Hz) as it had also shown pre-movement synchronization. However, unlike beta, alpha synchronization exhibited a less robust burst-like behavior, with less than 2 bursts per trial on average (Figure S4A, B, C). As a consequence, we characterized alpha dynamics across trials with the mean alpha power. This trial-wise averaged alpha was tested for its predictability of trajectory error evolution using the regARMA model. The optimal model selection procedure was repeated using the same steps carried out for beta bursts, which led to the choice of regARMA(1,1) as the one with the least AIC value for each perturbation (Table S3). Further, we tested the model using population leave-one-out cross-validation and compared the prediction errors of these models using alpha and beta bursts as external predictors. Here, we found that the model prediction error was consistently larger for the model using alpha mean power independent of the induced perturbation (N-PMSE; 2×2 repeated measures ANOVA; Main Effect of Model (beta bursts *vs.* mean alpha): F (2,32) = 5.04, p = 0.039, ^2^= 0.24, Figure S4D). Because we were forced to use for each trial mean alpha power (rather than alpha bursts), we performed a control analysis replacing beta thbursts with average beta power in each trial and found that the results remained similar (N-PMSE; 2×2 repeated measures ANOVA; Model (mean beta *vs.* mean alpha): F (2,32) = 6.84, p = 0.019, ηp = 0.30, Figure S4E).

## Discussion

Our findings provide a comprehensive understanding of the predictive processes underlying adaptative behavior. By integrating both conventional and dynamic analytical approaches, we offer novel insights into (i) how pre-movement beta activity emerges from the switch between different environments; and (ii) how outcome expectation is progressively updated when transitioning to stable brain states, enabling performance improvement and maintenance of successful behavioral outcomes. Our insights span three levels. Anatomically, by identifying cerebello-cortical networks as key in predictive processes supporting adaptive behavior; connectivity-wise, by showing directional and dynamical network interactions mediated by beta oscillations from cerebellar and parietal to prefrontal areas; and behaviorally, by establishing the trial-by-trial contribution of pre-movement beta burst activity to motor performance stabilization.

Using a conventional analysis of averaged beta activity, we first showed that changes in environmental characteristics impacted pre-movement beta activity. Pre-movement beta synchronization increased during stable states when the characteristic of the environment was known and motor errors were minimized, compared to the transition state, when a switch from one environment to another had just occurred. We argue that the increase in pre-movement beta synchronization occurs regardless of performance improvement based on two facts. First, beta power amplitude was equally low in the transition state for both perturbation conditions (rotation and no rotation) despite large differences in trajectory error magnitude. Second, pre-movement beta synchronization during stable states was more pronounced in the absence of rotation (during which motor performance did not improve) compared to when rotation was applied. This modulation of pre-movement beta synchronization up to 2.5 s before movement onset aligns with previous studies using transient and unpredictable perturbations (*10*, *11*) which show systematic attenuation of beta synchronization following trials with larger trajectory errors. With our design consisting of consecutive trials with the same perturbation, we elegantly demonstrate that high trajectory errors are not a necessary condition for a subsequent attenuation of pre-movement beta activity. Instead, it is the switch to a yet unknown environmental characteristic, leading to lower outcome predictability. The originality of our contribution lies in the investigation of pre-movement beta synchronization when, following the switch of environment, outcome expectation is built over several trials of an environment with the same characteristics. The early timing of this signal does not overlap with *prediction errors*, as pre-movement beta synchronization occurs before target appearance when motor plans, from which prediction errors are generated, are not yet available. We hypothesize that pre-movement beta activity encodes the build-up of an anticipatory signal reflecting the response of the sensorimotor system to the switch between environments with different characteristics. As participants build a representation of the environment and produce a more stable motor performance, beta power increases up to 2.5 seconds before movement onset, serving as an anticipatory signal informed by previous trials. Importantly, building such an anticipatory signal is facilitated in environments with no perturbation (no rotation) where motor performance is overall more stable and predictable.

Using MEG, we identified for the first time two distinct brain networks associated with beta oscillations: one identified in the building of expectations to stabilize motor performance when facing environmental changes, and another subtending the detection of specific environmental features while building expectations to maintain or stabilize motor performance. The first network included the posterior cerebellum bilaterally and the right temporal cortex. The cerebellum holds well-established credits in sensorimotor adaptation, with bilateral activations frequently observed during motor adaptation tasks (*50*). Conversely, the role of the temporal lobesin such a process remains debated (*29*). Even if not strictly necessary for the motor component of adaptation, the temporal cortex plays a significant role in the acquisition, implementation, and retrieval of action selection strategies. In this context, the concurrent involvement of the cerebellum and the temporal lobe may reflect their contribution to the formation of expectations useful to improve motor performance over trials. Instead, the second network included a broader set of regions, involving this time the right posterior cerebellum, the left parietal cortex, and the right prefrontal cortex. Two factors could possibly explain the asymmetry of cerebellar activity identified within the two networks. It may simply reflect statistical power: as participants performed the reaching task with their right hand, the right cerebellum, ipsilateral to the movement and contralateral to the motor cortices involved in task performance, was comparably more active than the left cerebellum. Alternatively, the right posterior cerebellum may be more involved in cognitive processes related to the formation of an anticipatory signal, i.e. outcome expectations (*50*, *51*), as compared to the left cerebellum (*52*, *53*). Interestingly, the right cerebellum has been shown to contribute to cognitive aspects of implicit motor sequence learning (*51*), its activity being modulated by environmental uncertainty. Predictive processes within the cerebellum allow building inference to account for uncertainty, as well as sensorimotor predictions on a trial-by-trial basis to estimate future motor states (*54*, *55*). We, therefore, hypothesize that this second network contributes to accumulating evidence to update the expectations based on environmental characteristics, reduce uncertainty, and stabilize motor outcomes (*56–58*). Unlike prediction errors, increased beta synchronization may consist of a general anticipatory signal that predicts the success (rather than failure) of expected events to occur. As these expectations are refined, humans can engage in metacognitive processes, evaluating errors from previous trials, and informing motor plans of the subsequent movement.

The effective connectivity further revealed how this information, supported by beta oscillations, flows in the cerebello-cortical network and unfolds under two different time scales. First, we observed a global bidirectional connectivity increase between the cerebellum, parietal, and prefrontal cortices from the transition to the stable state. Second, within a single trial, connectivity dynamically increased between the caudal regions (cerebellum, parietal) toward the prefrontal cortex in time windows closer to motor planning and target appearance. We hypothesize that these three areas coordinate operations to identify the characteristics of a set environment, by operating jointly, albeit with varying degrees of specialization. While the cerebellum and the parietal cortex are frequently implicated in predictive processes leading to adaptation (*27*), the involvement of the prefrontal cortex is generally associated with strategic processes of error correction (*35*). Our results suggest that the prefrontal cortex receives cumulative information from both the cerebellum and parietal cortex, potentially aiding in the implementation of appropriate strategies to favor the stability of motor outcomes.

Our finding associating anticipatory beta activity in the cerebellum due to changes in outcome expectations and enviromental characteristics has paramount implications for our field. First, for many decades the cerebellum was conceived as a mere motor structure, involved in the online correction of ongoing movements. In recent years, studies have suggested a role in processes occurring before movement execution, and event anticipation (*31–33*). Our study further supports the role of the cerebellum in cognitive aspects of adaptive behavior (*53*). Additionally, while beta activity within the cerebellum has been shown before (*15*, *19*, *21*), we bring an important brick of knowledge showing how this signal is timely used to progressively improve motor performance. Finally, localizing cerebellar sources has long remained controversial (*59*, *60*). Our MEG data repeatedly converge with independent data-driven analyses, reinforcing the reliability of our findings. This was likely facilitated by careful positioning of participants’ heads to optimize sensor caudal coverage, the use of individualized head models including the cerebellum, and time-frequency analyses capturing non-phase-locked activity, thus mitigating signal cancellation due to the complexity of cerebellar anatomy (*60*).

While the results from averaged beta synchronization highlighted the functional role of the cerebello-cortical network in generating an anticipatory signal tied to the expectations of environmental characteristics and motor outcome, trial-by-trial analyses offered complementary mechanistic insights into adaptive behavior. The evolution of beta bursts suggests the spontaneous emergence of order from what initially appears as a disorganized system in the early trials. According to complex system theory, the brain learns from experience through emergent activity, with brain states and behavior connected by non-linear relationships (*2*, *61*). Specifically, beta bursts likely originating from the collective interaction of brain components within the cerebello-cortical network may support the emergence of a stable state. Through predictive modeling of trial-by-trial changes in both motor performance and beta burst power, we provided, for the first time, evidence for the non-linear relationship between pre-movement beta bursts and motor performance. In the case of small perturbation (no rotation), beta burst amplitude explained the stability of behavior. In the case of large perturbation (rotation), beta burst amplitude contributed to explaining the drastic changes in behavior. Indeed, the brain-behavior relationship was strongest in blocks with larger trajectory errors (e.g., rotation; 25°–30°) compared to conditions with more stable error amplitudes (e.g., no rotation; 0°). These observations are not in contrast with the overall increase in pre-movement beta synchronization in blocks without perturbation. Instead, they suggest that pre-movement beta bursts encode information critical for anticipating trial-specific outcomes based on past experience. Trial-by-trial, the emergence of beta is an iterative process that supports the improvement and maintenance of motor outcomes in specific contexts. Although speculative, we propose that beta bursts may reflect the activity of an expanding pool of neurons responding more synchronously to increase the system stability, recruiting a set of structures within the network to fine-tune future motor plans. Alternatively, the progressive increase in beta burst amplitude could be related to action inhibition: as participants learn the rules governing the environment trial by trial, their knowledge of what to do may lead to increased eagerness to initiate the movement. However, we consider this unlikely, as action initiation is not linked to the mean beta burst power over a large time window (up to 2.5 seconds before movement onset). Instead, it is associated with the timing and amplitude of the final burst occurring closer to the movement onset (*44*, *45*). It is generally closer to movement onset that beta activity serves as an indicator of movement preparation, as final motor plans are enacted (*5*, *62–64*), and it is more closely related to whether or not to initiate a movement (*20*, *63*).

In conclusion, we provide important insights into the anatomical structures and coding brain mechanisms supporting the emergence of expectations during motor adaptation. We propose a unifying perspective in which outcome expectations and emergent properties of brain activity are tied to the same physiological assumption: that, following a change in the environment, beta oscillations serve as a neural anticipatory signal that allows recovery of behavioral stability. Our findings also open the field to a new conception of cerebellar function, with the idea that beta burst signals in cerebellar networks could perform the same anticipatory or predictive function across diverse domains, including language, spatial memory, reward learning (*65*). Our study, though, is also hindered by some current limitations that will be addressed in future work. First, the specific contributions of the cerebellum and the parietal cortex on this pre-movement anticipatory signal could not be disentangled. These regions may operate in tandem, but future work, using non-invasive brain stimulation could help causally delineate their distinctive roles. Second, while MEG improved the localization of cerebellar beta activity, its yet limited spatial resolution compared to functional Magnetic Resonance Imaging (fMRI), calls for caution when pinpointing the role of deep brain sources. Future work assessing changes of beta activity in other forms of learning, or clinical populations with cerebellar dysfunction (e.g., cerebellar ataxia) or diseases with altered beta activity (*e.g.*, Parkinson’s disease (*64*), essential tremor (*21*)) will shed further light on such questions and confirm their relevance in building up outcome expectations.

## Materials and Methods

### Population

We recruited 17 right-handed (as defined by the Edinburgh Questionnaire(*66*)) healthy volunteers (12 females, 45.7±14.1 years; mean ± std) with no history of medical, neurological, or psychiatric conditions, and with normal or corrected-to-normal vision. All participants met the additional MRI safety criteria (*67*) and provided written informed consent. The study was approved by the ethics committee “Ile de France VI - Groupe Hospitalier Pitié-Salpétrière” (INSERM C13-45) and conducted in accordance with the Declaration of Helsinki.

### Experimental paradigm and task description

Participants performed a modified version of a classic sensorimotor reaching task (*42*), consisting of blocks of 50 trials under either normal or rotated visual feedback. Participants controlled a joystick (Thetix, Current Designs, Philadelphia, USA) with their dominant right hand. A gray cursor on the screen indicated the joystick’s position. Upon target presentation, participants were instructed to move the cursor towards the visual target swiftly (taking the straightest path). In the normal visual feedback blocks, the cursor’s trajectory matched the joystick position and the expected visual feedback (0° rotation), while in the rotated visual feedback the trajectory was rotated either 25° clockwise (25° rotation) or 30° counterclockwise (−30° rotation) relative to the actual joystick trajectory. Rotation amplitudes were selected to induce noticeable trajectory errors and elicit explicit motor adjustments. Stimuli were displayed on a screen with a white background using MATLAB (The MathWorks, Natick, Massachusetts, United States of America (USA)) and its Psychophysics Toolbox. Each trial began with the presentation of a solid gray cursor (0.36° radius, viewed from 80 cm) within a 0.72° radius fixation circle at the bottom center of the screen. After 4.2 s (jittered by ±1.15 s) a circular black target (0.57° radius) appeared at one of five possible locations (9.3° radially from the fixation circle; Figure 1). Target locations were pseudorandomly determined, ensuring equal probability for each location across trials. Each trial ended when the cursor remained within the target for 0.3 s, causing the target to turn green and disappear (Figure 1). Participants were then instructed to return the cursor to the central fixation circle. The maximum reaching movement duration was 2.5 seconds; if participants exceeded this time limit, the target disappeared, prompting participants to return the joystick to the central location. All participants completed two separate experimental sessions on different days. The two sessions occurred at least one week apart but no more than 6 weeks apart. Both sessions were identical, each comprising six blocks of 50 trials of the reaching task, with two blocks per rotation (0°, 25°, or −30° rotation), for a total of 4 blocks per rotation. Blocks with different rotations were randomly interleaved to prevent retention effects across different sessions. The interleaved rotation blocks required participants to continuously update their sensorimotor predictions from one block to another, even in those blocks in which no rotation was applied (*42*).

### Behavioral data

We quantified ‘trajectory errors’ as the Area Under the Curve (AUC) representing the deviation of the actual trajectory from an ideal straight line between the start and target positions (in cm²). The start position was defined as the cursor’s location at the time point when it left the fixation circle for at least three consecutive time frames, whereas the endpoint was determined as the cursor’s position when it entered the target boundary. For trials in which the reach movement exceeded 2.5 seconds, resulting in missing data (0 to 6.5% of the total amount of trials, with an average of 3.6 % of trials for the whole population), we used interpolation to estimate these values. Specifically, we used a power law function to fit the collected data (*68*), defined as follows:

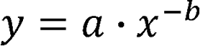

Where *y* is the AUC, *x* is the trial number, *a* is the scaling constant, and *b* is the exponent. We used nonlinear least squares fitting to estimate the parameters *a* and *b* from the observed data, and these coefficients were then used to interpolate missing AUC values. To differentiate between transition and stable states, we identified the trial where on average participants reached 80% of their error adjustments within a block (*69*). This threshold captured the transition point in which error reduction stabilized. To do so, for each participant, we calculated a power fit curve for the AUC values and used its first derivative to determine when 80% of the cumulative change had been achieved. On average, this corresponded to the first 16 trials within each block. Thus, the first 16 trials of each block included errors which were categorized as the transition state, whereas for consistency purposes, the last 16 trials of each block represented the stable state (Figure 2A). AUC values were then averaged across these states and blocks with similar perturbations (rotation and no rotation). We assessed motor performance improvement using a 2×2 repeated measures ANOVA with *State* (transition *vs.* stable) and *Perturbation* (rotation *vs.* no rotation) as within-subjects factors. The significance level was set at α=0.05, with the Huynh-Feldt correction applied when necessary to account for violations of sphericity. When appropriate, Bonferroni-corrected post-hoc comparisons were conducted.

### Brain data

#### T1-anatomical image acquisition and preprocessing

We first acquired individual T1-weighted MRI scans (MP-RAGE sequence) using an 8-channel head coil within a 3T MRI system (Verio system, Siemens, Germany) to enable accurate spatial co-registration with the MEG sensors and compute individual head models. The MRI scans were then segmented with Freesurfer (*70*), resulting in a cortical mesh with 15,000 vertices and a cerebellar mesh with 2,000 vertices (Brainstorm; (*71*)).

#### MEG data acquisition and preprocessing

Participants’ oscillatory brain activity was recorded using a 306-sensors whole-head MEG system (Megin Triux), comprising 102 radially oriented magnetometers and 204 horizontal gradiometers. Only gradiometers were used in the analysis due to their higher sensitivity to local cortical sources. Participants’ heads were carefully positioned to optimize sensor caudal coverage. Data were digitized at 1000 Hz with a band pass filter of 0-300 Hz. Fiducial markers and other anatomical boundaries were recorded with a 3D-Polhemus FASTRAK® digitizer, and vitamin patches were placed at the fiducial points for accurate registration with the MRI. Head movements were monitored using four coils positioned on the skull, while horizontal and vertical ocular movements were tracked through bipolar electrooculogram (EOG). Additionally, the electrocardiogram (ECG) was recorded using two electrodes placed on the left side of the abdomen and the right clavicle.

To reduce ambient noise, we used the temporal Signal Space Separation (tSSS) algorithm, a version of MaxFilter, in its standard configuration. Unlike the typical approach of segmenting the data into 20-second windows, the analysis was performed on the entire recording to avoid introducing discontinuities in the resulting signal. Further preprocessing was conducted using MATLAB and its open-source Fieldtrip toolbox ((*72*); www.fieldtriptoolbox.org). Specifically, the data was band-pass filtered (0.5–250 Hz, Butterworth filter), band-stop filtered (49.5–50.5 Hz) and low-pass filtered (40Hz). We then segmented the data into 5-second epochs centered around movement onset (from −3s to 2s). Trials containing artifacts such as SQUID jumps, eye blinks, head movements, and facial muscle activity, were excluded, leading to the removal of approximately 5.8% of trials.

#### Analysis of the average pre-movement beta activity in sensor space (MEG)

To analyze the average pre-movement beta synchronization, we transformed the MEG data into the time-frequency domain. We followed the Fieldtrip FLUX pipeline for low-frequency brain oscillations ((*73*); https://neuosc.com/flux/), consisting of a fast Fourier transformation for frequencies ranging from 2 to 40 Hz with a 500 ms sliding window using a single Hanning taper. The resulting power was then normalized relative to a common baseline (*7*, *11*). Specifically, power at each sensor, frequency, and time point was divided by the average power across all trials, independent of behavioral condition, and log-transformed. Time-frequency representations were, then, averaged across trials corresponding to the same behavioral state (transition and stable) and perturbations (0°, 25°, −30°). Data from the rotated visual feedback blocks (25° and −30°) were averaged, yielding four conditions per participant: rotation (transition and stable states) and no-rotation (transition and stable state). We performed statistical comparisons using a 2×2 non-parametric cluster-based approach (*74*) with *State* (transition *vs.* stable) and *Perturbation* (rotation *vs.* no rotation) as within-subject factors. We included all gradiometers (204) and individual time points between −2.5 s to 0 (relative to movement onset). We focused on the beta oscillations (14-26 Hz), avoiding potential overlap with the alpha range and aligning with prior studies associating beta modulation with sensorimotor adaptation (*11*, *12*).

To assess pre-movement beta synchronization changes across states and perturbations, we first computed differences between the transition and stable state within each perturbation condition (Δ(stable, transition)). We then run a two-tailed paired t-test at each sample to compare power changes between perturbations (Interaction: rotation, Δ(stable, transition) *vs.* no rotation Δ(stable, transition)). Adjacent time points and sensors were organized into clusters (threshold of p^Δ^ < 0.025). These resulting clusters were then evaluated for statistical significance against a permutation distribution, generated through 100,000 permutations involving random shuffling of conditions across all participants (significance level p < 0.025). We also ran a cluster analysis to test the main effects of *State* and *Perturbation* on pre-movement beta synchronization. To assess the main effect of *State*, we combined the data from both perturbations (rotation and no rotation) and statistically compared the two states (stable *vs.* transition). Similarly, to test the main effect of *Perturbation*, we combined the data from both states (transition and stable) for both perturbations and then statistically compared the two perturbations (rotation *vs.* no rotation). When appropriate, we performed post-hoc cluster tests using the same statistical approach.

#### Analysis of the average pre-movement beta activity in source space (MEG)

To localize pre-movement beta sources, participants’ MRIs and MEG sensor positions were first aligned using the fiducial landmarks and the scalp markers. The individual volume conduction models were then constructed using single-shell head model (*75*), derived from MRI segmentations. The lead field matrix was computed based on each participant’s source and volume conduction models, encompassing the cortex and cerebellum (3D grid 1 cm resolution). To identify sources of pre-movement beta activity indicated by the cluster analysis, we used the Dynamical Imaging of Coherent Sources (DICS) approach implemented in Fieldtrip (*76*). We opted for this approach because it is particularly suited for localizing sources in the time-frequency domain (*77*). Source estimates generated by beamforming approaches, including DICS, are prone to a noise bias toward the center of the head (*76*, *77*). This bias was corrected by contrasting conditions where possible (for example, Interaction: rotation, Δ(stable, transition) *vs.*no rotation, Δ(stable, transition)) or by applying the neural activity index ^Δ^AI; (*78*)) when direct contrasts were unavailable (for example, Main Effect of *State*: stable *vs.* transition). To estimate the power at each specific brain location (grid), we used participants’ lead fields and computed individual cross-spectral density (CSD) matrices. CSD matrices were computed using the multitaper method on the beta band (20 Hz, ±6 Hz, covering 14-26 Hz). We focused on sensors and time points identified as significant in the cluster analysis (*State* x *Perturbation* Interaction). Although cluster-based permutation tests on time-frequency representations do not directly establish the significance of specific sensor and time-latency differences (*79*), we could reasonably assume that the identified sensors and time points corresponded to regions where the effects were strongest. Finally, to highlight the specific regions contributing the most to pre-movement beta activity, we normalized the individual source signals using an MNI template with a regularly spaced (8 mm) three-dimensional grid of locations covering the brain volume. These normalized sources were then imported into SPM12 (Statistical Parametric Wellcome Department of Cognitive Neurology, London, UK), for further analysis. We performed paired t-tests comparing the relevant conditions identified by the sensor-space cluster-based results. Source locations were identified by identifying power peak-maxima (threshold: p < 0.001) to anatomical regions defined by the AAL atlas (*80*).

#### Dynamic connectivity analysis: nonparametric Granger causality (MEG)

We assessed the dynamic connectivity between relevant areas identified by the source analysis. Specifically, we generated a source model by specifying regions of interest (ROIs) using predefined MNI coordinates for parietal, frontal, and cerebellar as defined by the AAL atlas (left parietal inferior, right frontal inferior, and right Crus II; Table S1). To extract source signals, we applied a linearly constrained minimum variance (LCMV) beamformer, which is well-suited for localizing sources in the time domain (*23*). As we wanted to determine the temporal evolution of this connectivity, we analyzed five overlapping 0.45 s time windows before target appearance (0.1 s overlap; from −2.5 to −2.05; from −2.15 to −1.7; from −1.8 to −1.35; from −1.45 to −0.9; from −1.0 to −0.65). These time-windows were selected to capture at least six cycles of beta oscillations at the lowest frequency (14 Hz). For each grid location corresponding to the three ROIs, we computed the CSD matrix of source signals using a fast Fourier transform with 5 Hz frequency smoothing. The resulting power was then averaged across grid locations belonging to the same ROI. For the connectivity computation, we applied a multivariate nonparametric Granger causality approach. Specifically, we used a blockwise method to estimate directional influences between pairs of sources conditional on the remaining source (*81*). This analysis yielded six directional connectivity values (two for each pair of ROIs) across the five time-windows.

Finally, to assess how connectivity evolves during the pre-movement period, we performed six separate 5×2×2 repeated measures ANOVAs with *Time Window* (−2.5 to −2.05, −2.15 to −1.7, −1.8 to −1.35, −1.45 to −0.9, and −1.0 to −0.65 s), *State* (transition *vs.* stable), and *Perturbation* (rotation *vs.* no rotation) as within-subject factors. Significance was set at α = 0.05, with Huynh-Feldt corrections applied when necessary, and Bonferroni post-hoc tests conducted where applicable.

#### Analysis of the single trial-based pre-movement beta activity

To analyze pre-movement beta activity at the single-trial level, we adopted the approach by Shin et al. (*49*). We focused on sensors showing significant changes from transition to stable states when comparing the different perturbations. We analyzed a time window from −2.5 to 0 s relative to the movement onset, and frequencies between 2 and 40 Hz. The time-frequency transformation used a complex Morlet wavelet characterized by 7 wavelet cycles. The obtained power was normalized by calculating each frequency’s median power across each trial, separately for each participant and block (factors of median; FOM). Beta bursts were defined as regions within the beta range (14-26 Hz) where the power exceeded 3 times the median power (3X). We chose this cut-off because it yielded the best representation of the trial-by-trial variability in pre-movement mean power. We determined this threshold empirically as it provided the best representation of trial-by-trial variability in mean pre-movement beta power. Specifically, we quantified the percentage of the spectrogram area exceeding thresholds ranging from 1 to 7 times the median power and correlated this percentage with the mean beta power for each trial (Pearson’s test). The 3X threshold yielded the strongest correlation (please see Results; Figure 6B).

Beta bursts can be quantified in different ways. We, therefore, needed to choose the beta feature that best represented the process we aimed to characterize. To do this, we first selected the beta features identified by previous work as relevant to behavior (i.e., bursts mean power, burst number, last burst power; (*44*, *49*)). We then correlated these features with the average beta power (Pearson’s test) and used repeated measure ANOVA with beta features as a within-subject factor (beta bursts mean power *vs.* bursts number *vs.* last burst power) to identify the feature with the strongest correlation (i.e., events mean power; Figure 6D). Finally, to verify the relevance of the selected beta feature to motor adjustments, we compared the temporal dynamics of beta burst power to background beta activity. Background activity was defined as the power outside burst periods within each trial. For trials removed due to artifacts, missing values were estimated using a moving median interpolation (8-trial window). A 2×2×2 repeated measures ANOVA was conducted with *Feature* (beta burst power *vs.* background beta activity), *State* (transition *vs.* stable), and *Perturbation* (rotation *vs.* no rotation) as within-subject factors. For all ANOVAs, the significance was set at α = 0.05, with Huynh-Feldt correction applied when necessary, and Bonferroni post-hoc tests where applicable.

### Brain-behavior relationship: predictive modeling

#### Model framework

To quantify the relationship between participants’ trajectory error evolution and beta burst dynamics, we used a regARMA model, treating the data across trials as a time-series. The regARMA(p,q) model is given by the following equation:

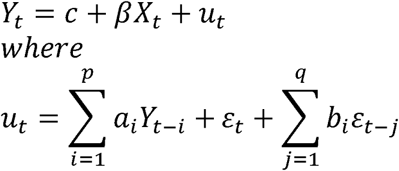

Where *Yt* is the dependent variable (trajectory error amplitude at trial t), *Xt* is the external predictor (beta burst power at trial *t*), β represents the influence of *Xt* on *Yt* (beta bursts contribution to AUC), *p* is the AR order, and *q* is the MA order.

Because regARMA model architecture assumes stationarity, before model implementation, we tested for stationarity of the time-series involved (*Y* and *X*). This analysis was performed across all perturbations on the average data across 17 participants, using the Augmented Dickey-Fuller test. Both trajectory errors and beta burst power time-series were found to be stationary (all p values < 0.05, Table S4).

#### Optimal model selection

To specify the regARMA model architecture, we determined the autoregressive (AR) order, moving average (MA) order, noise distribution (Gaussian *vs.* t-distribution), and intercept term. To select the optimal AR and MA orders we used the population average trajectory errors (*Yt*) and beta bursts mean power (*Xt*) across all participants. We examined the Autocorrelation Function (ACF) and Partial Autocorrelation Function (PACF) and noted the lag (m for ACF and n for PACF) at which these sharply dropped below two standard deviations of the data while using a trend stationary model. We then determined the optimal model architecture by comparing all feasible ARMA models by fixing AR and MA orders at n and m respectively, and varying noise distributions (Gaussian *vs.* t-distribution) and the inclusion of an intercept term. Model selection was based on the lowest AIC value among models where β significantly contributed to explaining variance. After fixing the noise distribution and deciding on the inclusion or exclusion of an intercept term based on the AIC values, we also tested models with AR and MA orders increasing and decreasing from (n,m) to check if the AIC values improved. A summary of AIC values for all feasible tested ARMA models is provided in Table S2. We selected regARMA (1,1) as the optimal candidate because it was consistently the best model for all rotations with the least AIC values. Only rotation −30° had two possible candidates, i.e., ARMA (1,1) and ARMA (1,2). However, as the difference between their AIC values was lower than 2, we opted for the model consistent with the other rotations.

#### Model parameter estimation and validation

We validated the optimal model selected across the population of 17 participants using a leave-one-out approach. For each participant and perturbation, we estimated the model parameters using data from the population average of the remaining 16 participants. The stationarity was reconfirmed for each of the averages calculated from the different combinations of 16/17 participants before the parameter estimation. These parameters were then used to predict the left-out participant’s trajectory error time-series. Specifically, we used the participant’s error from the first trial, along with the trial-by-trial beta burst dynamics over the 50 trials of a block, to predict the trajectory error across the remaining 49 trials (Figures 7A; see Figures S1, S2 and S3 for individual simulations). To quantify prediction errors, we calculated the prediction mean squared error (PMSE) between the model’s predictions and observed data, from the 2nd to 50th trials.

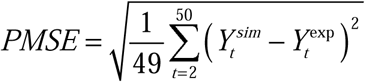

Given that PMSE can vary with the scale of the data (e.g., trajectory errors amplitude at 0° rotation are typically smaller than those at 25° or −30°), we min-max normalized the PMSE (N-PMSE) to ensure fair comparisons across different conditions, e.g., perturbations or different models. In the min-max normalization was done as

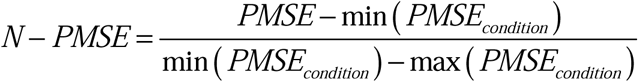

To test whether beta burst power differently influenced error amplitude evolution across perturbations, we conducted repeated-measures ANOVA with *Perturbation* (0° *vs.* 25° *vs.* −30°) as a within-subject factor using beta contribution (β) as the dependent variable. Significance was set at α = 0.05, with Huynh-Feldt corrections β and Bonferroni post-hoc tests applied where necessary. Similar analyses were conducted to compare the model prediction accuracy (N-PMSE) across perturbations.

#### Model specificity

We also tested whether the relationship between trajectory errors and beta burst power was specific to frequency. To this aim, we replaced beta burst power with alpha mean power while predicting reach errors. Alpha mean power was used instead of alpha burst power due to the absence of reliable pre-movement alpha bursts in our data (Figure S5). We employed the same model selection, estimation, and validation processes described in the previous section (for details on alpha mean power time-series stationarity and model selection, please see tables S4 and S5). To determine the specificity of the original model, we compared model fit (AIC) using a 2×3 repeated-measures ANOVA with *Model* (beta bursts *vs.* alpha mean power) and *Perturbations* (0° *vs.* 25° *vs.* −30°) as within-subject factors. Significance was set at α = 0.05, with Huynh-Feldt corrections applied when necessary, and Bonferroni corrected post-hoc tests conducted where applicable. The same approach was used for a control analysis, where we compared models using mean alpha power with those using mean beta power as external predictors (for details on alpha and beta mean power time-series stationarity and model selection, please see tables S3, S5 and S6).

## Acknowledgments

This work was carried out in the MRI and MEG platform of the CENIR within the Paris Brain Institute (ICM).

## Fondation pour la Recherche Médicale (CGa)

European Union’s Horizon 2020 research and innovation program; the Marie Słodowska-Curie grant agreement 897941 (MB)

## Authors contributions

Conceptualization: MB, SM, DS, TP, CG. Methodology: MB, VV, CGa. Data Collection: QW, AR, CGa. Data Analysis: MB, VV, VR, QW, MR. Visualization: MB, VV, CGa. Supervision: AVC, DS, TP, CGa. Writing—original draft: MB, VV, CGa. Writing— review & editing: MB, VV, VR, WQ, MR, AR, CGi, SM, AVC, DS, TP, CGa.

## Data and materials availability

The data will be publicly available upon publication. Data analysis was performed using publicly available software and toolboxes as described in the Methods section. Any additional information may be requested from the corresponding authors (MB, CG).

## Supplementary Materials

Supplementary Materials contain:

Figures S1 to S4

Tables S1 to S6

