## Supplemental Figures and Tables for "Emergence of Anticipatory Beta Activity to Facilitate Behavioral Stability Following Environmental Changes"

Martina Bracco *et al.*

**This PDF file includes:**

Figures S1 to S4
Tables S1 to S6

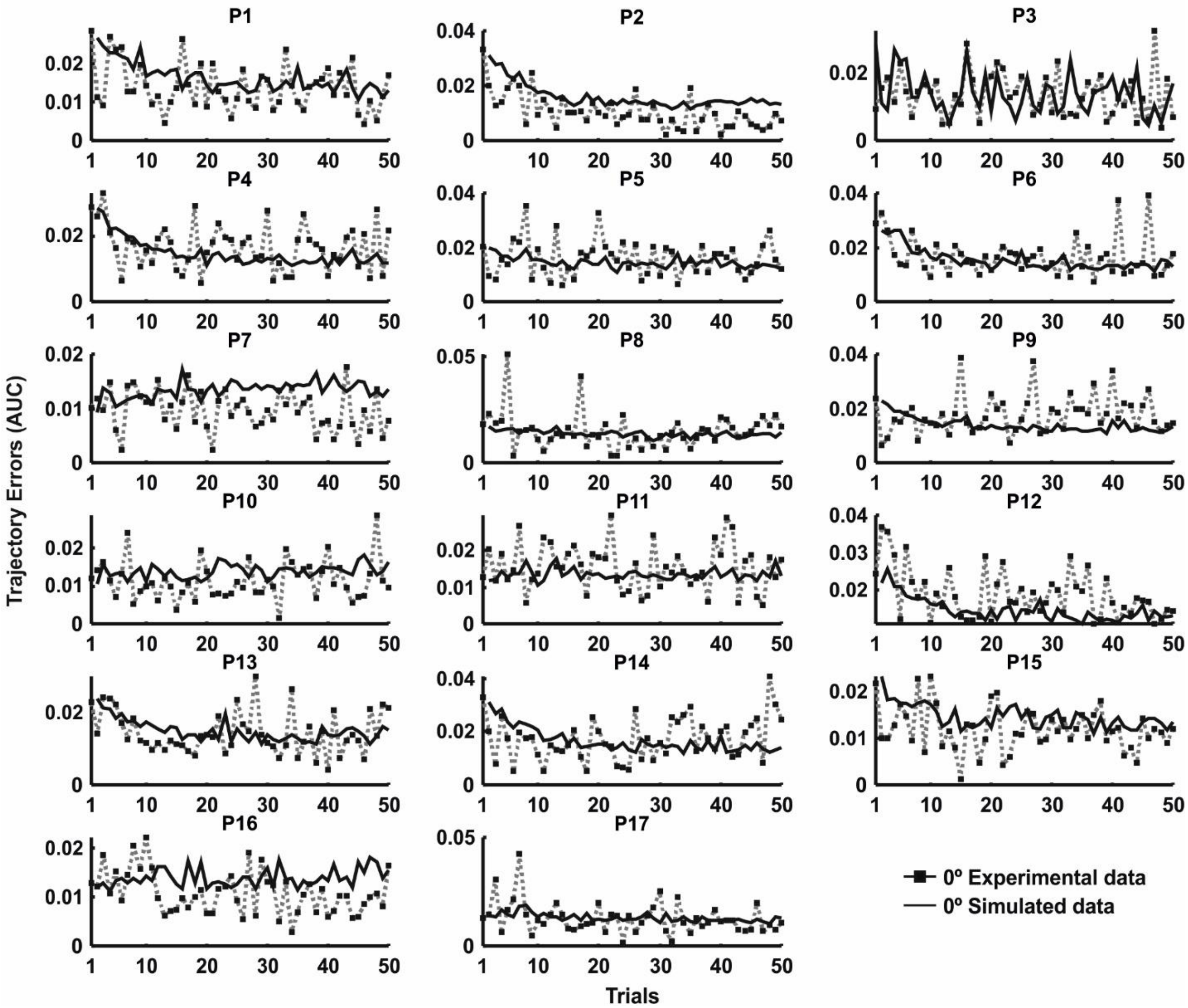


**Fig. S1. Predictive model results on single participants for 0° perturbation.**

Real and predicted trajectory errors along 50 consecutive trials, averaged across 0° perturbation blocks, for each of the 17 participants. Real trajectory errors are quantified as the area under the curve (AUC), representing the deviation from the ideal straight-line trajectory between the start and target positions. Higher AUC values indicate greater trajectory errors. Predicted errors are estimated by validating the chosen regARMA(1,1) model using a leave-one-out cross-validation approach. Each plot corresponds to a different participant.

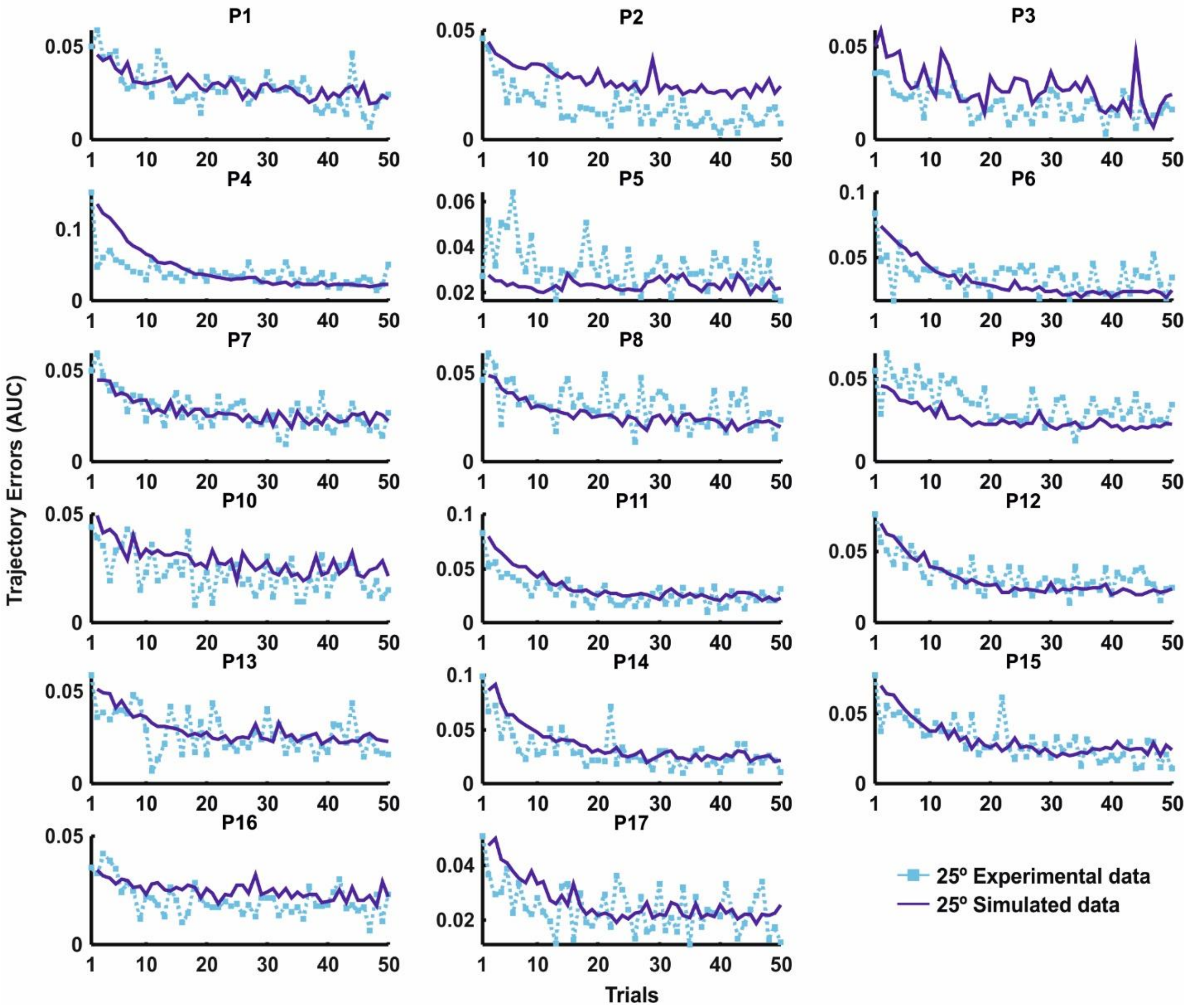


**Fig. S2. Predictive model results on single participants for 25° perturbation.**

Real and predicted trajectory errors along 50 consecutive trials, averaged across 25° perturbation blocks, for each of the 17 participants. Real trajectory errors are quantified as the area under the curve (AUC), representing the deviation from the ideal straight-line trajectory between the start and target positions. Higher AUC values indicate greater trajectory errors. Predicted errors are estimated by validating the chosen regARMA(1,1) model using a leave-one-out cross-validation approach. Each plot corresponds to a different participant.

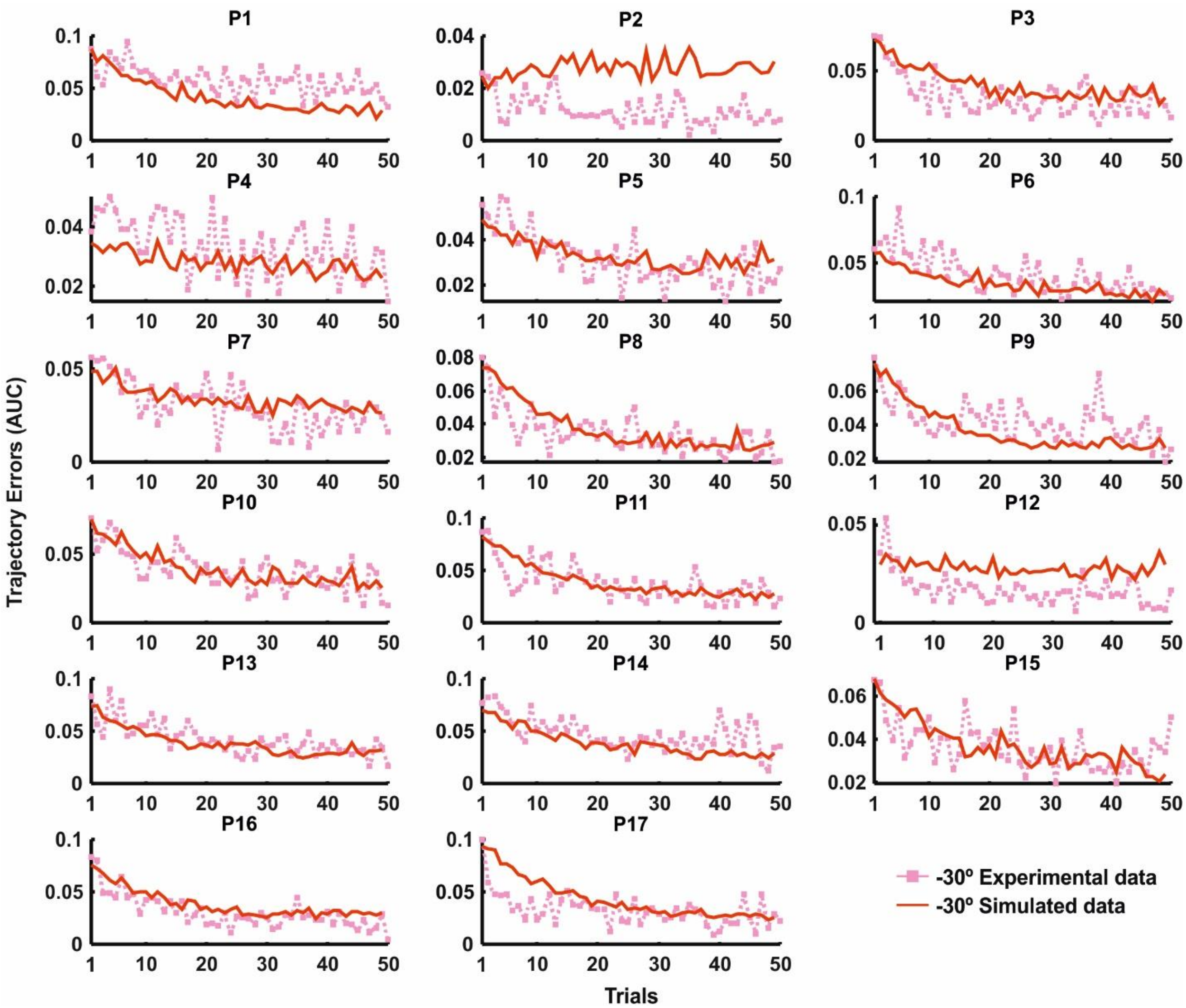


**Fig. S3. Predictive model results on single participants for -30° perturbation.**

Real and predicted trajectory errors along 50 consecutive trials, averaged across -30° perturbation blocks, for each of the 17 participants. Real trajectory errors are quantified as the area under the curve (AUC), representing the deviation from the ideal straight-line trajectory between the start and target positions. Higher AUC values indicate greater trajectory errors. Predicted errors are estimated by validating the chosen regARMA(1,1) model using a leave-one-out cross-validation approach. Each plot corresponds to a different participant.

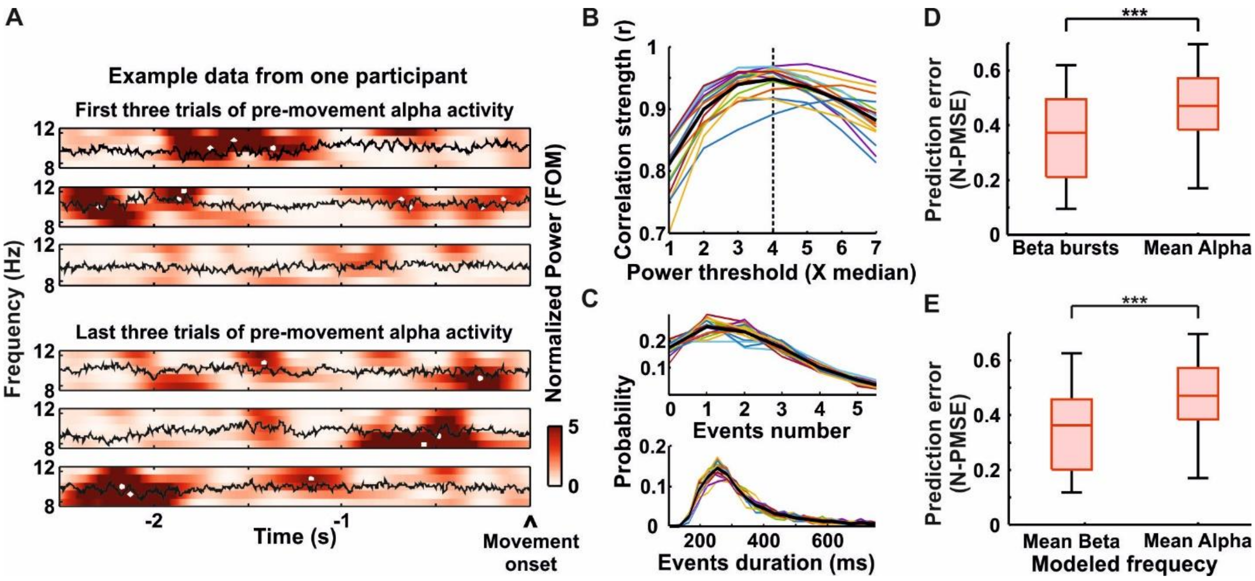


**Fig. S4. Alpha (8-12 Hz) bursts identification and mean alpha power modeling results.**

**(**A) Alpha bursts in the 8–12 Hz range are shown from -2.5 seconds to movement onset for the first three and last three trials of an example block in a representative participant. Data were normalized as factors of the median (FOM) for each frequency, calculated separately for each participant and condition. (**B**) The threshold for data normalization was selected based on the highest correlation between the percentage of the spectrogram area exceeding thresholds ranging from 1 to 7 times the median power and the mean beta power for each trial. Colored lines represent the averages of trials and blocks for individual participants, while the black line shows the group average. The dashed vertical black line indicates the threshold with the highest correlation (4X median power). (**C**) Probability distribution of events number (top) and events duration (bottom), where an event is an alpha burst exceeding the 4X median cutoff. Distributions are plotted as the average across trials and blocks for individual participants (colored lines), with the population average in black. (**D**) Results of the 2x2 repeated measures ANOVA comparing the normalized prediction mean squared error (N-PMSE) of the regARMA(1,1) models across different models (Main Effect of Model: beta bursts vs mean alpha). (**E**) Results of the 2x2 repeated measures ANOVA comparing the normalized prediction mean squared error (N-PMSE) of the regARMA models across different models (Main Effect of Model: mean beta vs mean alpha). Box plots show the median (horizontal line), the lower and upper quartiles (box), the minimum and maximum values that are not outliers (whiskers), and the outliers, computed using the interquartile range (dots). *** indicates $p < 0.001$

| Anatomical Position (AAL Atlas) | MNI Coordinates (mm) x | y | z | t-Values (peak) | p-Values (peak) |
|---|---|---|---|---|---|
| **MAIN EFFECT of Error Stage** | | | | | |
| Cerebellum_Crus1_L | -56 | -48 | -38 | 6.73 | p < 0.001 |
| Cerebellum_Crus1_L | -14 | -88 | -32 | 4.80 | p < 0.001 |
| Insula_R | 36 | 12 | -16 | 4.54 | p < 0.001 |
| Cerebelum_Crus2_R | 50 | -72 | -42 | 4.14 | p < 0.001 |
| **INTERACTION (Error Stage * Error context)** | | | | | |
| Parietal_Inf_L | -28 | -58 | 38 | 8.04 | p < 0.001 |
| Frontal_Inf_Tri_R | 56 | 32 | 22 | 7.06 | p < 0.001 |
| Paracentral_Lobule_L | 14 | -94 | 34 | 4.76 | p < 0.001 |
| Parietal_Sup_R | 22 | -64 | 50 | 4.35 | p < 0.001 |
| Paracentral_Lobule_R | 16 | 40 | 50 | 3.89 | p = 0.001 |
| Cerebelum_Crus2_R | 34 | -74 | -44 | 3.79 | p = 0.001 |

**Table S1. Source analysis results.**

Results of the paired t-tests in source space comparing the relevant conditions identified by the sensor-space cluster-based results. This analysis aims at highlighting the specific regions contributing the most to pre-movement beta activity. Source locations were identified by identifying power peak-maxima (threshold: $p < 0.001$) to anatomical regions defined by the AAL atlas. Only peaks identified within the brain space and belonging to the AAL atlas are reported.

| ARMA order (p, q) | Distribution of noise term (ε) | Intercept term | AIC |
|---|---|---|---|
| **Rotation 0°** | | | |
| (1,1) | Gaussian distribution | Absent | - 497.01* |
| (1,0) | | | - 468.22 |
| (1,1) | t- distribution | Absent | - 495.02 |
| **Rotation 25°** | | | |
| (1,1) | Gaussian distribution | Absent | - 421.34* |
| (1,0) | | | -379.16 |
| (1,1) | t- distribution | Absent | - 419.32 |
| (1,0) | | | -377.11 |
| (2,0) | | | -389.47 |
| **Rotation -30°** | | | |
| (1,1) | Gaussian distribution | Absent | - 423.94* |
| (1,0) | | | - 404.89 |
| (1,2) | | | - 425.55 |
| (1,1) | t- distribution | Absent | - 421.93 |
| (1,0) | | | - 402.82 |
| (1,2) | | | - 423.55 |

**Table S2. Possible regARMA model architectures using trajectory errors (AUC) time series as the dependent variable and beta-bursts power times series as the external predictor.**

Asterisks indicate the selected optimal model from all possible combinations for each rotation. The model was selected based on the lowest Akaike Information Criterion (AIC). The maximum AR and MA ders were determined by examining the PACF and ACF respectively, and noting the lag (n and m) at which they dropped below two standard deviations of the data with the use of a trend stationary model. Only model architectures showing significance for AR term(s), MA term(s), beta and variance are shown.

| ARMA order (p, q) | Distribution of noise term (ε) | Intercept term | AIC |
|---|---|---|---|
| **Rotation 0°** | | | |
| (1,1) | Gaussian | Present | -492.04* |
| (1,0) | | Absent | -475.21 |
| (1,1) | t- distribution | Absent | - 491.74 |
| (1,0) | | | -471.52 |
| (0,1) | | | -457.56 |
| **Rotation 25°** | | | |
| (1,1) | Gaussian distribution | Absent | - 417.06* |
| (1,1) | t- distribution | Absent | -415.04 |
| **Rotation -30°** | | | |
| (1,1) | Gaussian distribution | Absent | - 423.32* |
| (1,0) | | | - 403.71 |
| (0,1) | | | - 326.50 |
| (1,1) | t- distribution | Absent | - 421.29 |
| (1,0) | | | - 401.62 |

**Table S3. Possible regARMA model architectures using trajectory errors (AUC) time series as the dependent variable and alpha mean power times series as the external predictor.**

Asterisks indicate the selected optimal model from all possible combinations for each rotation. The model was selected based on the lowest Akaike Information Criterion (AIC). The maximum AR and MA orders were determined by examining the PACF and ACF respectively, and noting the lag (n and m) at which they dropped below two standard deviations of the data with the use of a trend stationary model. Only model architectures showing significant AR term(s), MA term(s), beta and variance are shown.

| Rotation | 0° | | 25° | | 30° | |
|---|---|---|---|---|---|---|
| | AUC | BB | AUC | BB | AUC | BB |
| ADF stationarity test p values | 0.001 | 0.001 | 0.006 | 0.022 | 0.010 | 0.005 |

**Table S4. Stationarity tests on reach errors (AUC) and beta-burst (BB) power times series for all rotations.**

To determine stationarity, we noted the lag (n) at which PACF dropped below two standard deviations of the data with the use of a trend stationary model. We determined the lag with the lowest Akaike Information Criterion (AIC) between 0 and m in the ADF test and the corresponding p-value at that lag. P values $< 0.05$ suggest stationarity.

| **Rotation** | **0°** | | **25°** | | **30°** | |
|---|---|---|---|---|---|---|
| | **MA** | **MB** | **MA** | **MB** | **MA** | **MB** |
| **ADF stationarity test p values** | 0.022 | 0.001 | 0.008 | 0.004 | 0.035 | 0.009 |

**Table S5. Stationarity tests on mean beta (MB) and alpha (MA) power times series for all rotations.**

To determine stationarity for mean beta power, we noted the lag (n) at which PACF dropped below two standard deviations of the data with the use of a trend stationary model. For mean alpha, we noted the lag at which PACF drops below three standard deviations of the data. For both, we determined the lag with the lowest Akaike Information Criterion (AIC) between 0 and m in the ADF test and the corresponding p-value at that lag. P values $< 0.05$ suggest stationarity.

| ARMA order (p, q) | Distribution of noise term (ε) | Intercept term | AIC |
|---|---|---|---|
| **Rotation 0°** | | | |
| (1,1) | Gaussian distribution | Absent | -500.74* |
| (1,0) | | | -473.57 |
| (0,1) | | | -456.93 |
| (2,0) | | | -478.94 |
| (1,1) | t- distribution | Absent | -498.75 |
| (0,1) | | | -463.26 |
| **Rotation 25°** | | | |
| (1,1) | Gaussian distribution | Present | -382.84 |
| (1,1) | | Absent | -421.19* |
| (1,0) | | | -378.13 |
| (1,1) | t- distribution | Absent | -419.20 |
| (1,0) | | | -376.09 |
| **Rotation -30°** | | | |
| (1,1) | Gaussian distribution | Absent | -425.90 |
| (1,0) | | | -411.02 |
| (1,2) | | | -429.02* |
| (1,1) | t- distribution | Absent | -424.23 |
| (1,0) | | | -408.97 |
| (1,2) | | | -427.43 |

**Table S6. Possible regARMA model architectures using trajectory errors (AUC) time series as the dependent variable and beta mean power times series as the external predictor.**

Asterisks indicate the selected optimal model from all possible combinations for each rotation. The model was selected based on the lowest Akaike Information Criterion (AIC). The maximum AR and MA orders were determined by examining the PACF and ACF respectively, and noting the lag (n and m) at which they dropped below two standard deviations of the data with the use of a trend stationary model. Only model architectures showing significant AR term(s), MA term(s), beta and variance are shown.
